# DYRK1A inhibition by the postbiotic metabolite Urolithin A contributes to enhanced β-cell function: Structural, Biochemical and Cellular Evidence

**DOI:** 10.64898/2026.11.23.768402

**Authors:** Michał Nowacki, Alicja Bis, Katarzyna Pustelny, Ali Maruf, Mayurdhvaj Parbatsinh Rathod, Alex Matsuda, Przemysław Grygier, Jakub Piwowarski, Anna Czarna, Maciej Dawidowski

## Abstract

Urolithin A (UA), a postbiotic metabolite of ellagitannins (ETs), has shown potential in modulating metabolic disorders but the mechanisms underlying these activities remain incompletely understood. Here, we investigate the influence of UA on β-cell function and assess the role of dual-specificity tyrosine-regulated kinase 1A (DYRK1A) inhibition in mediating these effects. We develop a novel transition-metal-free synthesis of UA employing a one-pot triple Baeyer-Villiger oxidation. Biophysical and biochemical analyses demonstrate that UA, and to a lesser extent its glucuronides (UA 3/8-gluc), bind to and inhibit DYRK1A, whereas the dietary precursor ellagic acid (EA) is inactive. Cellular studies confirm the intracellular DYRK1A engagement by UA but not by UA 3/8-gluc. To elucidate the molecular basis of this inhibition, we solve the crystal structure of UA-DYRK1A complex, providing the first high-resolution structural data on UA binding to a protein target and explaining the differential activities of UA, UA 3/8-gluc, and EA. Studies in MIN6 cells show that UA enhance glucose-stimulated insulin secretion (GSIS) and increase cell proliferation to levels comparable with the potent DYRK1A inhibitor harmine. These findings propose a new mechanistic link between postbiotic metabolites of ETs and insulin homeostasis and establish a structural basis for development of urolithin-based kinase inhibitors.

## 1. Introduction

Gut microbiota-derived metabolites originating from dietary constituents have attracted increasing attention because of their diverse biological activities and their potential contribution to the maintenance of human health and disease modulation^1, 2^. Among them, metabolites of ellagitannins (ETs), polyphenolic compounds abundant in foods such as pomegranate, strawberries, raspberries, blackberries, and walnuts are of particular interest^3, 4^. Consumption of ET-rich food has been associated with a range of beneficial metabolic effects. For instance, pomegranate juice ingestion has been reported to improve selected parameters of glucose homeostasis in individuals with diabetes mellitus (DM), including reduction on fasting glucose and insulin resistance, and improvement in indices of pancreatic β-cell function^5^.

Importantly, ETs and their hydrolysis product ellagic acid (EA) exhibit very low oral bioavailability, making it difficult to directly attribute the biological effects of ET-rich food to these compounds themselves^4, 6^. However, it has been shown that ETs and EA are extensively metabolized by the human gut microbiota into more readily absorbed, biologically active urolithins (dibenzo[*b*,*d*]pyran-6-ones) and this discovery has provided a new perspective for understanding the biological effects of ET-rich foods^3^.

Among the systemically absorbable metabolites of ETs, urolithin A (UA) is one of the most extensively studied and has shown a broad range of biological activities in experimental models and, increasingly, in clinical studies^3, 7, 8^. Particular attention has been paid to its effects on mitochondrial homeostasis and muscle performance^9, 10^ which have been linked to the regulation of autophagy and mitophagy. In addition, urolithins have been reported to modulate metabolic processes and may be beneficial in DM and its complications, however the molecular mechanisms underlying these effects remain incompletely understood^11^. Previous studies indicated that UA, and, to a greater extent, other, structurally related ET metabolite - urolithin C (UC), enhances glucose-stimulated insulin secretion (GSIS) in β-cell models^12^. Mechanistic studies focused on UC showed that the compound increases GSIS through the activation of L-type calcium channels^12^ and ERK signalling^13^. In contrast, the molecular mechanisms of UA contributing to the similar effects remain undefined.

In parallel, the dual-specificity tyrosine-regulated kinase 1A (DYRK1A), has emerged as a promising molecular target in DM. Interference with DYRK1A activity by a range of chemical compounds, including the prototypical inhibitor harmine and other natural products, has been shown to stimulate β-cell proliferation in both rodent and human experimental systems and to improve functional parameters of β-cells^14–18^. Importantly, recent study has identified UA as an ATP-competitive inhibitor of DYRK1A in a cell-free biochemical assay and investigated this activity in context of neuroprotection^19^. However, whether UA can engage DYRK1A in living cells and whether inhibition of this kinase contributes to the reported effects of UA on pancreatic β-cell function has not been established.

Although UA has been demonstrated to exhibit a range of *in vivo* activities^8^, an important aspect when considering the clinical potential of the compound is its low plasma exposure. Although the compound is readily absorbed in the gut, it simultaneously undergoes extensive phase II conjugation. Consequently, urolithins are found in serum and tissues predominantly in the form of *O*-glucuronides^20^. Moreover, distinct urolithin metabotypes exist, defined by the profile of urolithins produced by an individual gut microbiota^21, 22^. These metabotypes may vary with factors such as ageing and diet, and, importantly, not all individuals are able to produce the same urolithins, including those associated with the most pronounced biological activities. Another challenge is that, although the microbial conversion of EAs to urolithins can be efficient, it may be difficult to consume sufficient amounts of ellagitannin-rich foods on a regular basis to achieve biologically relevant concentrations of urolithins in body fluids and tissues^3^. Taking these limitations into account, and considering extensive phase II conjugation of UA, the development of dietary supplements containing defined, high does of synthetic compounds has gained increasing attention in recent years. UA has attained GRAS status in the United States^23^, and health effects of supplementation with UA have already been evaluated in several clinical trials^9, 10^. Despite this growing demand, the chemistry of UA remains underdeveloped and only few strategies for its chemical synthesis are reported in the scientific literature. Moreover, due to the wide spectrum of reported biological activities of UA, the compound represents an attractive starting point for structure-guided medicinal chemistry campaigns and for development of molecular probes. Such efforts, however, may be hampered by lack of experimentally determined, high-resolution structural information on the compound binding mode to its target protein(s). Although computational models of UA binding several proposed targets have been reported^19, 24–26^, a crystallographically validated binding model is required to define the molecular determinates of recognition and to guide rational modification of the scaffold.

Considering the aforementioned, important gaps in the knowledge of mechanisms by which UA and ET-rich foods impact metabolic disorders, in this study we investigate DYRK1A inhibition as a potential mechanism contributing to the effects of UA on β-cell function. Further, we investigate whether UA and its metabolites engage the kinase in living cells and define the precise structural basis of these interactions by solving the co-crystal structure of UA with DYRK1A. This first, experimentally determined crystal structure of UA bound to its protein target, together with a new synthetic route to the compound, provides a structural foundation for the rational design of urolithin-based derivatives with improved biological properties.

## 2. Results

### 2.1. First metal-free synthesis of UA based on a one-pot, triple Baeyer-Villiger oxidation/hydrolysis sequence

Despite the therapeutic potential of UA and its increasingly widespread use in dietary supplements in relatively high daily doses (e.g. 1000 mg), only few synthetic strategies towards this compound have been published in the scientific literature so far. The commonly employed methods are based on the intermolecular condensation (**Figure S1**, path A) of resorcinol and 2-halobenzoic acids in the presence of a base and copper catalyst (Hurtley reaction)^27^ ^28, 29^; or on an intramolecular lactonization either by the AcOH-triggered cyclization (Figure S1, path A) between a tertiary amide and a phenol^30^ or by the Cu(I)-mediated C-O_carboxylate_ bond formation (Figure S1, path B) in subcritical water^31^. These strategies have certain drawbacks. For example, they require introduction of appropriate protection of OH groups, which, in turn, necessitates their subsequent cleavage. Furthermore, transition metal compounds, specifically copper or palladium, are typically employed in the key C-C coupling reactions in these approaches. With an aim to expand the synthetic repertoire towards UA we sought for alternative strategies for the central lactone ring formation from alternative starting materials, without introduction of protecting groups in additional steps and avoiding transition metal catalysis.

A well-established protocol utilized the Baeyer-Villiger oxidation to convert fluorenone to 6*H*-benzo[*c*]chromen-6-one^32^, which is a core scaffold in UA. Inspired by this, we decided to explore whether the lactone formation and the simultaneous, exocyclic 3,8-bishydroxylation leading to UA can occur in the-one pot protocol, in the course of a triple Baeyer-Villiger oxidation of a suitable tris-ketone (*i*.*e*., 3,8-bisacetoxyfluorenone)(**Figure 1A**).

**Figure 1.**
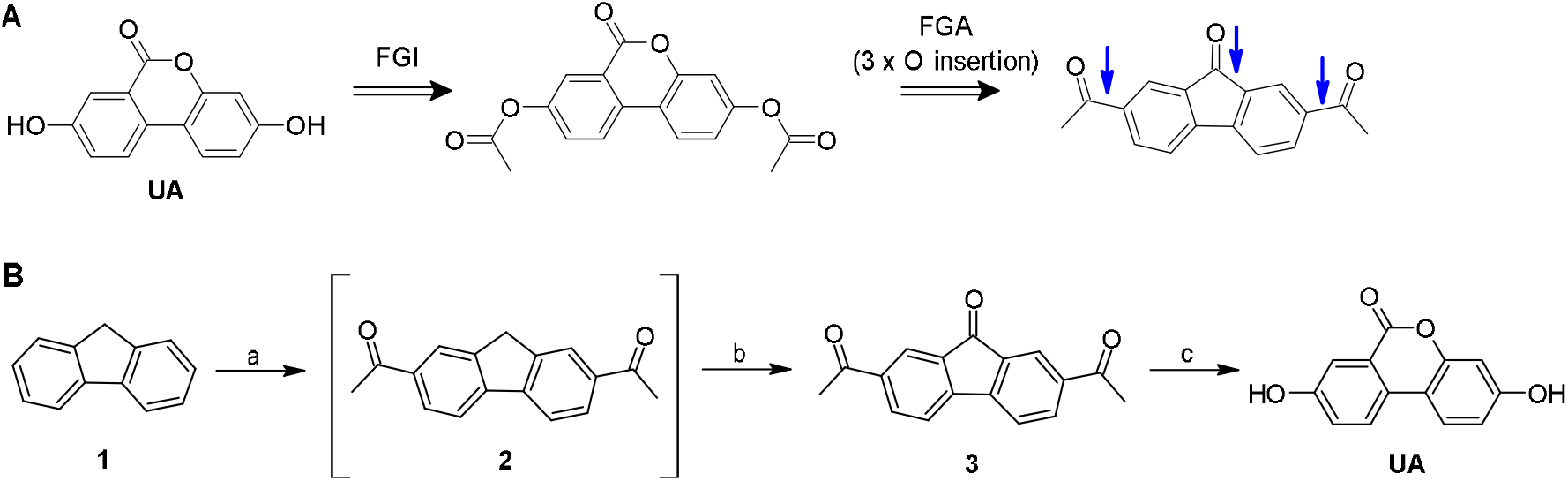
Synthesis of UA. A) The retrosynthetic strategy to UA based on triple Baeyer-Villiger oxidation/hydrolysis. FGI - Functional Group Interconversion; FGA - Functional Group Addition. B) Reagents and conditions. a) Ac_2_O (4 equiv.), AlCl_3_ (8 equiv.), DCM, - 10°C – rt, 3 h; then b) O_2_ (air), DMF, K_2_CO_3_ (10 mol%), 30°C, 4 days; overall yield 72%. c) 1. TFA (35 equiv.), H_2_O_2_ 30% (15 equiv.), 5°C, 64 h; 2. NaHCO_3_, H_2_O; 0°C; then 1M HCl_aq_ / MeOH (1:1), 70°C, 4 h; overall yield 39%.

The developed metal- and protecting group-free synthetic pathway to UA is presented in **Figure 1B**. Readily available, cheap fluorene 1 was bis acetylated using Ac_2_O *via* Friedel-Crafts acylation^33^. The literature reports the oxidation of the resulting bis-ketone 2 to the tris-ketone 3 with K_2_Cr_2_O_7_ in AcOH^34^. However, in our hands, despite the additional purification rounds, product 3 retained Cr(III) species. To circumvent this issue, and, therefore, to ultimately accomplish a first, fully transition-metal-free synthesis of UA, we successfully applied alternative oxidation methods of crude 2 employing deprotonation of the mildly acidic methylene bridge in the presence of catalytic amount of base and the subsequent oxidation of the generated carbanion with molecular oxygen involving the single electron transfer (SET) mechanism^35^. The developed, simple protocol consisted in stirring the crude acetylation product 2 with 10 mol% of K_2_CO_3_ in DMF for 4 days on air to give intermediate 3 in 72% yield (from 1, over 2 steps).

UA was obtained from intermediate 3 by a one-pot sequence of 1) triple Baeyer-Villiger oxidation to simultaneously form a central lactone ring and introduce two exocyclic oxygen atoms; and 2) double hydrolysis to deacetylate the phenolic groups of UA. In the oxidation step, TFA was successfully used as a solvent and a precursor of one of the most powerful peroxy acids, which is formed *in situ* by the self-catalyzed esterification-type reaction with 30% aqueous solution of hydrogen peroxide. Importantly, the use of *m*-CPBA, the most common Baeyer-Villiger oxidant, led exclusively to the oxidation of *exo*-carbonyl groups (see: Supplementary Information). Attempts to further simplify the synthesis of UA by a Baeyer-Villiger oxidation of unprotected 3,8-dihydroxyfluorenone led to the full recovery of the starting material.

### 2.2. UA and its glucuronides, but not the primary precursor EA, bind and inhibit DYRK1A in a cell-free setting

Previous report provided the first evidence linking the biological activities of UA to its potent inhibition of DYRK1A^19^. Here, we examined whether DYRK1A inhibition is specific to UA or is also observed for compounds associated with its metabolic pathway, *i*.*e*., the dietary precursor EA and the mixture of II phase metabolites, UA 3- and 8-glucuronides (UA 3/8-gluc)^36^, for their ability to inhibit the recombinantly produced DYRK1A kinase domain in a cell-free setting.

The inhibitory potential of each compound was first assessed using the ADP-Glo kinase biochemical assay at a single compound concentration, using harmine, a well-studied DYRK1A inhibitor, as a positive reference (**Figure 2A**). Because UA is an ATP-competitive inhibitor of DYRK1A^19^, the assay was performed using ATP and the synthetic peptide substrate DYRKtide at their respective (K_M_) concentrations. This setup provided suitable conditions for detecting ATP-competitive inhibition and for comparing the relative potency of the tested compounds. The results showed that EA itself did not inhibit DYRK1A activity at the tested concentration of 20 μM, indicating that inhibitory activity emerges only after its metabolic conversion to UA and is retained in its glucuronidated metabolites. To quantitatively compare the inhibitory activity of UA and UA 3/8-gluc, we determined their (IC_50_) values against DYRK1A (**Figure 2B**). The glucuronide metabolites were approximately tenfold less potent than UA (∼20 μM vs 2.3 μM, respectively), however they still produced measurable inhibition of DYRK1A, showing that glucuronidation reduces rather than eliminates the activity.

**Figure 2.**
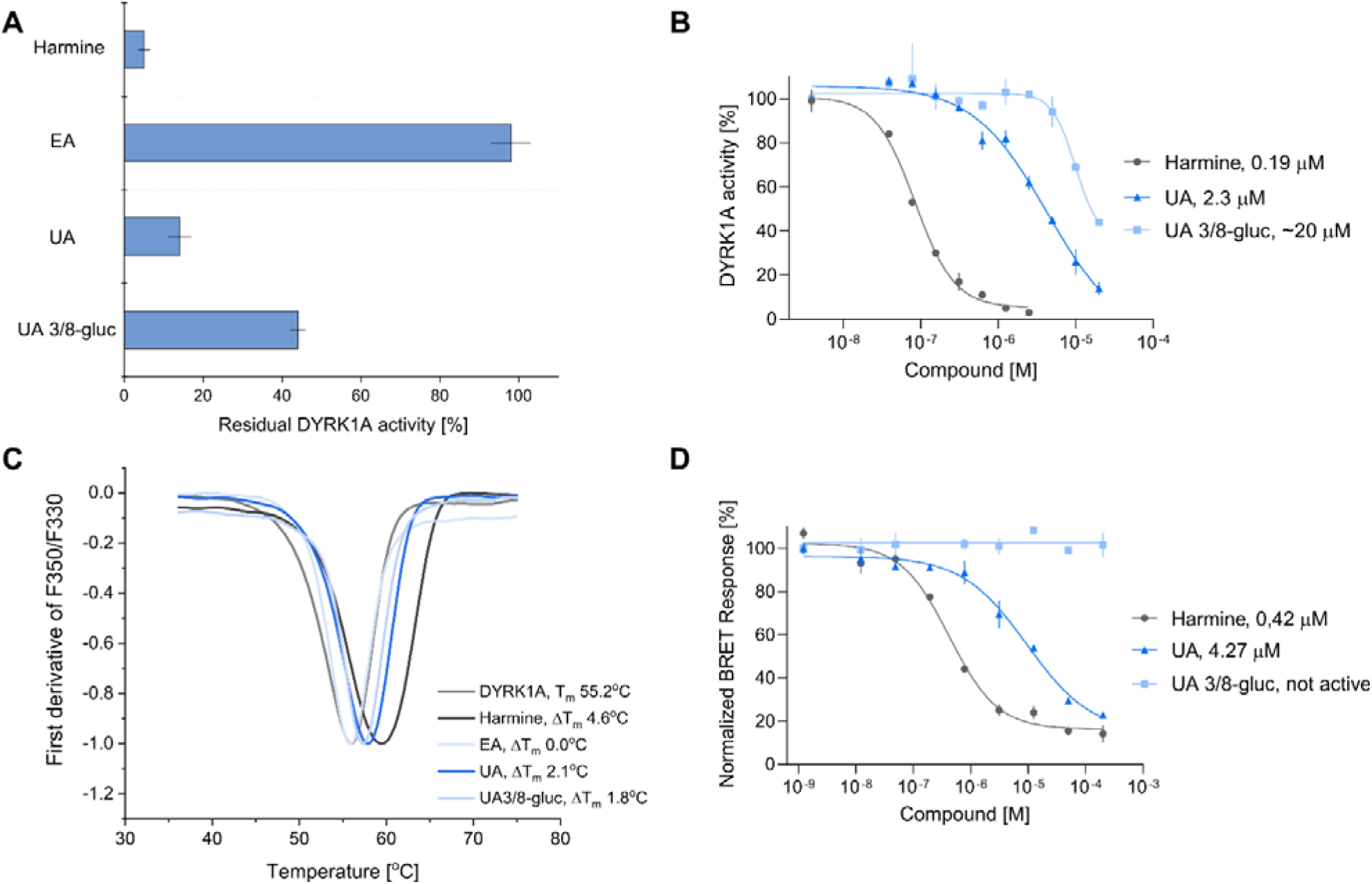
Effects of EA, UA, and UA 3/8-gluc on DYRK1A activity, thermal stability, and cellular TE. A) Residual DYRK1A activity in the presence of 20 µM harmine, EA, UA, and UA 3/8-gluc: harmine, 5 ± 1.5%; EA, 98 ± 5%; UA, 14 ± 3%; and UA 3/8-gluc, 44 ± 2%. B) Concentration–response analysis of harmine, UA, and UA 3/8-gluc using an *in vitro* peptide-based DYRK1A kinase assay. IC values were: harmine, 0.19 ± 0.019 µM; UA, 2.30 ± 0.16 µM; and UA 3/8-gluc, 20 ± 4 µM. C) nanoDSF analysis of compound-induced changes in DYRK1A thermal stability. Representative thermal unfolding curves from three independent measurements are shown. Melting temperatures were highly reproducible between experiments, with inter-experimental variation of ≤0.5°C. D) Evaluation of DYRK1A binding by harmine, UA, and UA 3/8-gluc in live cells using the NanoBRET TE assay. Cellular IC values were 0.42 ± 0.09 µM for harmine and 4.27 ± 0.77 µM for UA. All quantitative data are presented as mean ± SD from three independent experiments (n = 3).

To validate these biochemical observations, we performed a biophysical characterization using a nano differential scanning fluorimetry (nanoDSF) assay (**Figure 2C**), which enables precise determination of protein thermal stability and its modulation by the ligand. A positive shift in T serves as a direct indicator of protein stabilization upon ligand binding. The DYRK1A kinase domain alone exhibited a relatively high melting temperature (T 55.2°C), indicating intrinsic structural stability and reflects a well-engineered construct. Consistent with our enzymatic data, EA did not affect the thermal stability of the protein, confirming the lack of direct binding. In contrast, the reference inhibitor harmine markedly stabilized DYRK1A, increasing the T to 59.8°C. Both UA and the UA 3/8-gluc mixture also induced a clear thermal stabilization, with T values of 57.3°C and 57.0°C, respectively. These results demonstrate that, unlike the inactive dietary precursor EA, UA and its glucuronides directly interact with and stabilize the DYRK1A kinase domain.

### 2.3. Differential intracellular target engagement of DYRK1A by UA and its glucuronidated derivatives

While ADP-Glo assay and thermal stability measurement provide essential information on ligand affinity for the isolated DYRK1A kinase domain, they represent a simplified model that does not fully capture the complexity of the intracellular environment^37^. Therefore, we next performed nano bioluminescence resonance energy transfer (NanoBRET) target engagement (TE) assays in HEK293 cells, which enables real-time quantification of compound binding to full-length DYRK1A in living cells through displacement of a fluorescent tracer from a NanoLuc–DYRK1A fusion protein (**Figure 2D**). This assay allows to evaluate the inhibitory activity of a compound under physiologically relevant conditions, including high endogenous ATP concentrations and in the presence of cellular modulators, while providing insight into compound membrane permeability at the same time. The results revealed a clear distinction between the intracellular TE of active metabolite UA and its glucuronides. UA demonstrated a robust TE, with a cellular IC of 4.27 μM. This represents a moderate (∼twofold) shift compared to its biochemical IC (2.3 μM), indicating that UA effectively penetrates the cell membrane and retains its affinity for the DYRK1A catalytic site even in the presence of millimolar intracellular ATP concentrations. An analogous, ∼twofold shift was observed for harmine, included as a reference inhibitor, indicating that the cellular assay reliably captures the ATP-competitive behaviour under intracellular conditions. In contrast, UA 3/8-gluc was inactive in the cellular NanoBRET assay. Although these metabolites exhibited a measurable inhibitory activity in the ADP-Glo assay (IC ≈ 20 μM) and induced protein stabilization in nanoDSF experiments, they failed to displace the tracer in living cells, even at highest concentration tested (200 μM). This discrepancy highlights the substantial impact of phase II metabolism on the pharmacological profile of urolithins. While glucuronidation of UA only partially reduced intrinsic enzyme inhibition, it appears to limit cellular activity, most likely by impairing membrane permeability. Consequently, despite the predominance of phase II metabolites in circulation, the non-conjugated UA is the primary mediator of DYRK1A inhibition at the cellular level.

### 2.4. Structural basis of DYRK1A Inhibition by UA - a crystallographic depiction of binding to the ATP pocket

UA exerts its biological effects through interference with variety of metabolic pathways, including those mediated by protein kinases^38^. However, the precise molecular background of these activities remains poorly understood, and, despite UA has been suggested to bind several protein targets, the current knowledge on these interactions relies exclusively on molecular modelling^19, 24–26^. To date, no high-resolution co-crystal structures of UA bound to its putative protein targets have been reported that would experimentally verify the predicted binding models and serve as a guidance for medicinal chemistry and chemical biology. To address this important gap, we determined the first crystal structure of UA bound to its molecular target, DYRK1A.

The ligand is drawn with magenta bonds, and protein residues forming hydrogen bonds are drawn with grey bonds (carbon in black, oxygen in red, nitrogen in blue). Hydrogen bonds are shown as dashed lines with distances in Å, and water molecules are shown as dark red spheres. The direct hydrogen bonds are C8-OH – N (Leu241, 2.83 Å) and C3-OH – NZ (Lys188, 2.74 Å). The water-mediated interactions involve C8-OH - water (2.74 Å), linked to the Ser242 main-chain oxygen (2.97 Å) and the Leu241 carbonyl oxygen (3.19 Å), and C6-CO –water (3.32 Å), linked to the Asn244 side-chain amide (2.93 Å). Hydrophobic and van der Waals contacts are shown as spoked arcs and involve Ile165, Val173, Ala186, Phe238, Glu239, Met240, Leu294, Val306 and Asp307. In this diagram, protein side-chain interactions are shown only for contacts bridged by a single water molecule. Contacts mediated by longer water chains (e.g. Glu291) are therefore not shown, but they are visible in figure 3B. The figure was prepared with UCSC ChimeraX 1.10.1^39^.

**Figure 3.**
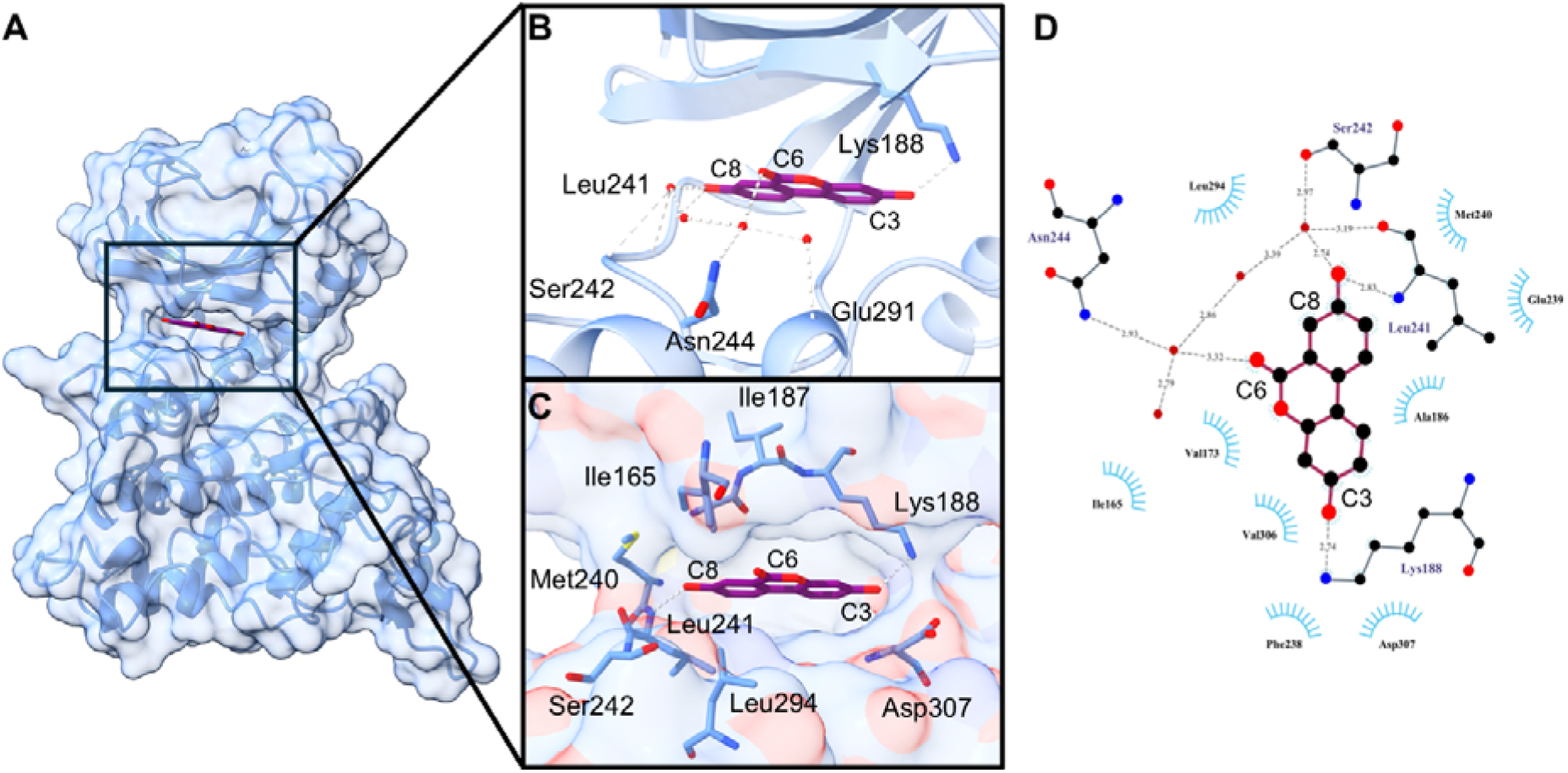
Binding mode of UA in the ATP-binding site of DYRK1A. A) Overall structure of the DYRK1A kinase domain (light blue) in complex with UA (magenta sticks)(PDB accession code: 35BW). UA is bound in the ATP-binding pocket. B) Detailed view of the polar interactions of UA (magenta sticks) with the protein (light blue). The C8 hydroxyl group forms a direct hydrogen bond (gray dashed lines) with the backbone amide of the hinge residue Leu241, and the C-3 hydroxyl group interacts directly with the side chain of the catalytic Lys188. The remaining contacts are mediated by a network of ordered water molecules (red spheres) linking the ligand to Ser242, Asn244 and Glu291. C) Close-up of the ATP-binding pocket, shown with the protein surface. The residues that contact the ligand directly or through water molecules (Lys188, Leu241, Ser242, Asn244 and Glu291) are shown. D) Two-dimensional schematic of the interactions of UA generated with LigPlot.

The DYRK1A kinase domain in complex with UA crystallized in the P2 2 2 space group, with the asymmetric unit comprising four molecules (**Table S1**). The crystals diffracted to a resolution of 2.13 Å. UA was clearly resolved in the ATP-binding pocket of DYRK1A, located in the cleft between the N- and C-terminal lobes (**Figure 3A**). Clear, well-defined electron density was observed for UA in all four protein molecules within the asymmetric unit (**Figure S2**), allowing its unambiguous placement in the active site. The four protein molecules adopted highly similar conformations, and the UA binding mode was conserved across all copies. Therefore, only molecule A was used for the subsequent structural analysis. The experimentally determined binding mode of UA to DYRK1A active site differs substantially from the previously published pose based on molecular modelling^19^. Compared with our data, the docking model shows the ligand adopting an inverted and twisted orientation, with the C3 hydroxyl group predicted to form a hydrogen bond with the backbone carbonyl of Glu239, and the C6 carbonyl group interacting with the backbone amide NH of Asp307. In contrast, in the high-resolution co-crystal structure, UA is oriented toward the hinge region of the ATP-binding pocket (**Figure 3B-D**). The C8 hydroxyl group of UA forms a direct hydrogen bond with the backbone of Leu241 (2.83 Å), anchoring the ligand within the binding site. At the opposite end of the tricyclic scaffold, the C3 hydroxyl group forms a polar contact with the catalytic Lys188 (2.74 Å). In addition, UA engages in an extensive network of water-mediated hydrogen bonds. Its C6 lactone carbonyl participates in a water-mediated interaction, forming a hydrogen bond with a nearby water molecule at a distance of 3.32 Å (**Figure 3B**). Although relatively weak, this interaction is geometrically plausible. The same water molecule participates in a broader network of polar contacts involving Ser242, Asn244, and Glu291, thereby complementing the direct hinge interactions and contributing to ligand stabilization within the active site. The inhibitor is further stabilized by hydrophobic contacts with Ile165, Val173, Ala186, Phe238, Met240, Leu294 and Val306, which form a complementary environment around the ligand (**Figure 3C**).

The incapability of EA to bind and inhibit DYRK1A in biochemical assays appeared intriguing, since both EA and UA are high-affinity ligands of the structurally related protein kinase CK2 (CK2)^28, 41, 42^. Importantly, the comparison of the obtained crystal data on UA-DYRK1A complex with the previously published EA-CK2 co-crystal structure (PDB accession code: 2ZJW) ^42^ provides a molecular rationale for this structure–activity relationship (SAR) (**Figure 4**). Based on this comparison, inability of EA to inhibit DYRK1A is unlikely to result from a general failure of EA to enter its ATP-binding pocket. Potent inhibition of CK2 by EA indicates that its planar, tetracyclic scaffold is fully compatible with kinase binding, but only when supported by an appropriate shape and interaction complementarity of the binding pocket. CK2 and DYRK1A share bulky phenylalanine gatekeeper residues, Phe113 and Phe238, respectively, which limit access to the hydrophobic back pocket. Consequently, Phe113 in CK2 constrains the extended EA scaffold entirely within the adenine-binding region. This spatial restriction prevents EA from making direct contacts with the hinge region. Instead, EA anchors to the CK2 hinge by a highly specific, stabilizing network of water molecules engaging Glu114 and Val116 backbones (**Figure 4B**). Similar interactions are possible with equally oriented Glu 239 and Leu241, respectively, in DYRK1A. Importantly, water-mediated binding of EA to CK2 occurs through His160 side chain in the upper part of the CK2 binding pocket. On the contrary, in DYRK1A this residue is mutated to Glu291, which rules out this important stabilization. The unique, water-mediated positioning enables EA to form critical, direct, polar contacts with side chains of two conserved residues in the catalytic Lys68 and Asp175 of the DWG motif in CK2. Strictly homologous Lys188 and Asp307 residues, respectively are present in DYRK1A and structural superimpositions suggest that EA could be anchored in its ATP-binding site in a similar fashion. Other, important nuances dictating selectivity of EA towards CK2, lie in the hydrophobic sandwich enclosing the ligand (**Figure 4C**). In CK2, this highly complementary enclosure relies heavily on residues Leu45, Val66 and Met163^42, 43^. The ligand-dependent conformational flexibility of Met163 may contribute to the ability of CK2 to accommodate the extended planar scaffold of EA by adjusting the lower hydrophobic boundary of the ATP-binding pocket. Together with Leu45, Met163 defines the depth and inclination of the plane occupied by flat aromatic ligands, thereby promoting complementary hydrophobic packing within the binding site (**Figure 4D**, top). In contrast, the corresponding hydrophobic residues in DYRK1A (Ile165 and Leu294) establish a much more rigid and constrained spatial environment (**Figure 4D**, bottom), creating steric clashes and preventing the rigid EA scaffold from simultaneously achieving the favourable hydrophobic enclosure and the polar interactions with the kinase. Thus, the contrasting activities of EA toward CK2 and DYRK1A likely reflect these kinase-specific differences in pocket complementarity and ligand stabilization. The critical role of this adaptable hydrophobic pocket in CK2 is further underscored by the reported mutational studies: introducing the Val66Ala and Met163Leu (V66A/M163L) double mutation in CK2 inclines the purine base binding plane, which not only dramatically shifts the enzyme’s co-substrate preference toward ATP over GTP, but also highlights how these bulky residues govern the binding affinity and orientation of planar inhibitors^43, 44^.

**Figure 4.**
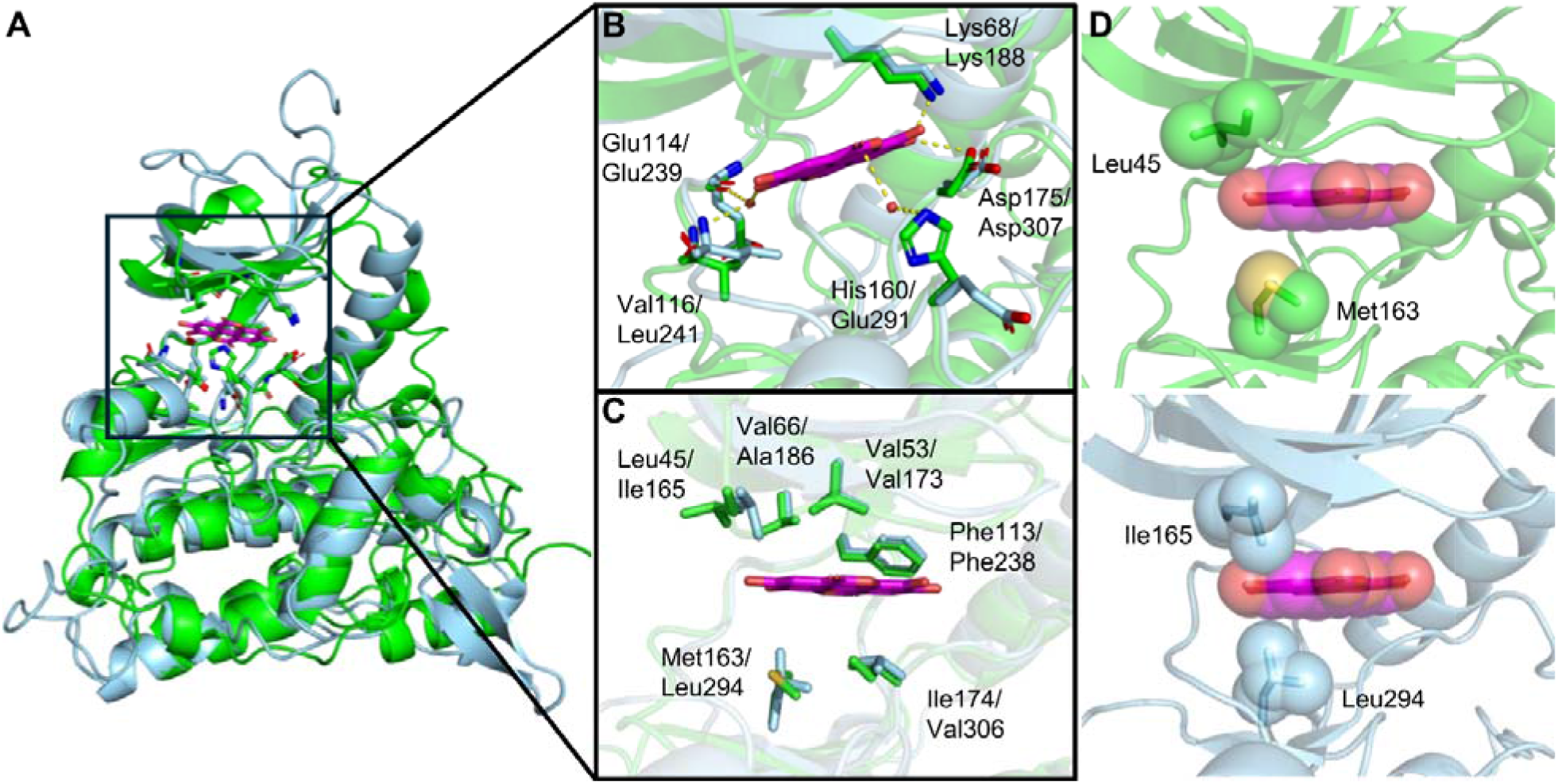
Kinase selectivity determinants of EA. A) Overlay of the EA-CK2 co-crystal structure (magenta and green, respectively, PDB accession code: 2ZJW) with DYRK1A (light blue, derived from PDB accession code: 35BW). B) Detailed view of selected, polar interactions (yellow dashes) of EA (magenta) with residues (Lys68, Glu114, Val116, His160, Asp175) in the active site of CK2 (green). CK2 is overlayed with DYRK1A (light blue). The respective Lys188, Glu239, Leu241, Glu191, Asp307 are shown. The His160 residue of CK2 which forms an important, water mediated contact with EA, corresponds to Glu291 in DYRK1A. C) Detailed view of hydrophobic enclosure of EA (magenta) in CK2 active site (green). CK2 is overlayed with DYRK1A (light blue). The selected, hydrophobic residues of both kinases are shown. D) The main differences in the spatial constraints of CK2 (top, green) and DYRK1A (bottom, light blue) active sites. The spheres represent vdW radii. The steric clashes occur between EA (red and magenta spheres) and Ile165 and Leu294 (light blue spheres) of DYRK1A. The figure was prepared with PyMOL 2.5.7^40^.

UA-DYRK1A cocrystal structure explains the weaker binding of UA 3/8-gluc, as compared to the parent compound. The bulky glucuronide substituents are likely to disrupt the key polar interactions formed by the hydroxyl groups of UA (Figure 3BD) and introduce steric constraints (Figure 3C), thus lowering the binding affinity. This not only explains the SAR of UA-related compounds studied herein but suggests that significant losses of affinity can be expected when using either of the UA hydroxyl groups as chemically tractable exit vectors in the kinase-oriented ligand design.

### 2.5. DYRK1A inhibition by UA contributes to enhanced β-cell function

The ability of UA to inhibit DYRK1A has previously been investigated in the context of neurodegeneration^19^, but the relevance of this activity to pancreatic β-cell function has not been explored. Other studies indicated that EA-derived metabolites, particularly UA and UC, may exert beneficial effects on β-cell function^12, 13^. However, mechanistic investigation focused exclusively on UC, showing that it enhances GSIS through the activation of L-type calcium channels^12^ and ERK1/2 pathway^13^. In parallel, DYRK1A has emerged as a potential therapeutic target in DM, as its inhibition can promote β-cell proliferation and support insulin secretory function^14, 15, 17, 18^. Having established that UA direct binds and inhibits DYRK1A *in vitro* and engages the kinase in cells, we next examined whether this activity could contribute to its previously observed effects on β-cells.

The cytotoxicity profile of UA in MIN6 cells demonstrated good biocompatibility across all tested concentrations (from 1 to 25 µM), with cell viability consistently remaining above 80%, indicating no harmful effects (**Figure 5A**). Interestingly, at lower concentrations (1 and 2.5 µM), UA slightly increased metabolic activity, as reflected by viability values exceeding 100% (111% and 107%, respectively). This enhancement may suggest a stimulatory effect on cellular metabolic activity or mitochondrial function, which is consistent with the known role of UA as a mitophagy inducer that promotes mitochondrial quality control^8, 45^. These findings are in line with previous reports demonstrating the safety profile of UA in various cell types (*e.g*., neuroblasts: 0.2-50 µM^45^, macrophages:10-50 µM^46^, and fibroblasts: 10-50 µM^47^) and support its suitability for further functional studies in pancreatic β-cells. Overall, the absence of cytotoxicity in MIN6 cells ensures that any observed effects on insulin secretion or cell proliferation are unlikely to be confounded by reduced cell viability.

**Figure 5.**
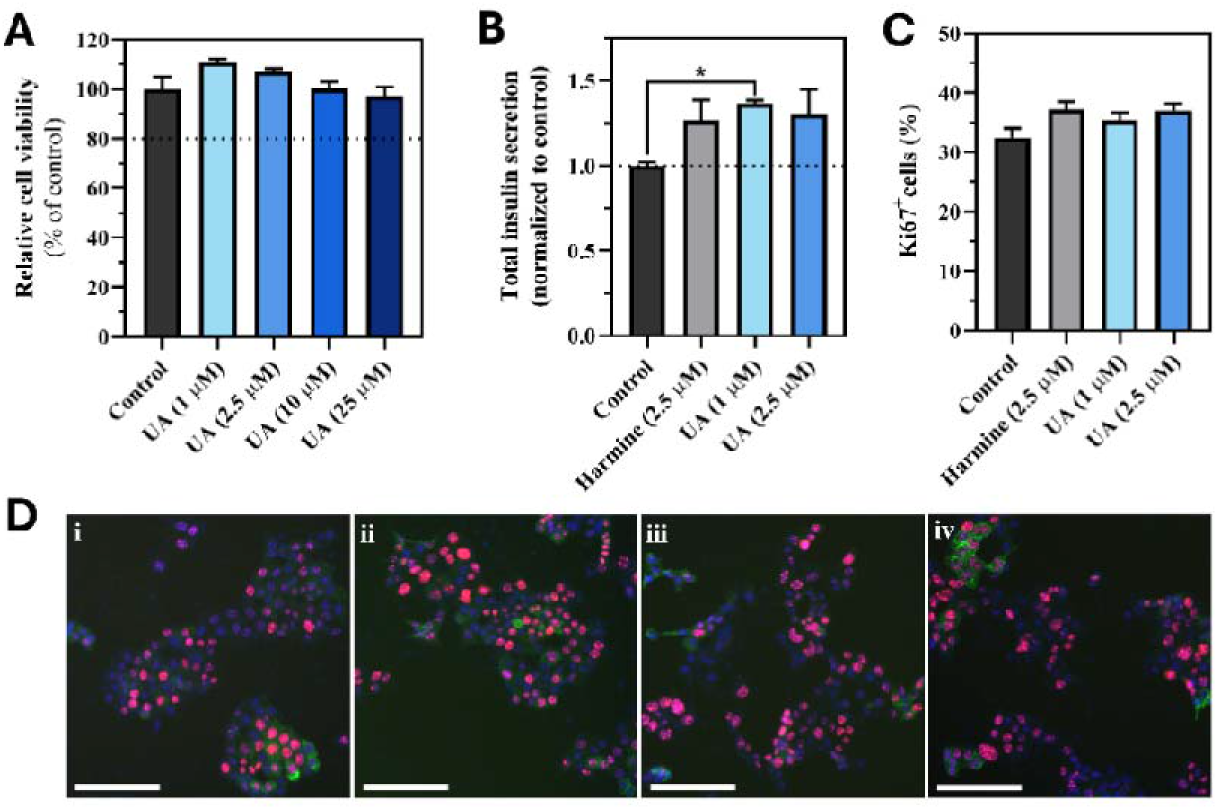
Biological effects of UA in MIN6 cells. A) Cell viability of MIN6 cells following treatment with UA at 1, 2.5, 10, and 25 µM for 72 h, assessed using the Alamar Blue assay. B) Total insulin secretion from MIN6 cells treated with UA at 1 and 2.5 µM and harmine at 2.5 µM for 72 h, measured by ELISA. C) Cell proliferation of MIN6 cells quantified as the percentage of Ki67+ cells after treatment with UA at 1 and 2.5 µM and harmine at 2.5 µM for 72 h. D) Representative fluorescence microscopy images corresponding to C): i) control, ii) harmine (2.5 µM), iii) UA (1 µM), and iv) UA (2.5 µM), respectively. DAPI: blue, Ki67: red, insulin: green. Scale bar: 200 µm. Data are presented as mean ± SEM (n = 3). Statistical significance was determined using One-way ANOVA followed by Bonferroni’s multiple comparisons test (*p < 0.05).

Low concentrations of UA significantly enhanced insulin secretion in MIN6 cells (**Figure 5B**). At 1 and 2.5 µM, the compound significantly enhanced insulin secretion (1.37- and 1.31-fold *vs.* control), surpassing harmine (1.27-fold), a well-characterized DYRK1A inhibitor used as a positive control. Furthermore, UA demonstrated a modest effect in MIN6 cell proliferation, as assessed by the percentage of Ki67 cells (**Figure 5CD**). Ki67 is a nuclear protein expressed during all active phases of the cell cycle (G1, S, G2, and M), but absent in quiescent cells (G0), and is therefore widely used as a marker of cellular proliferation^48^. At low concentrations (1 and 2.5 µM), UA slightly increased proliferation (35.4 and 37.0, respectively) compared to control (32.5%), reaching levels comparable to harmine (37.2%), a compound known to induce β-cell proliferation. Notably, this pro-proliferating effect coincided with the same low-µM range (1–2.5 µM) in which UA also enhanced insulin secretion and metabolic activity, suggesting these low concentrations represent a favorable functional window rather than a generic dose-response. Overall, these findings support a mild pro-proliferative effect of UA in MIN6 cells at the tested concentrations.

## 3. Discussion

Recently, it was demonstrated that UA is a potent, ATP-competitive DYRK1A inhibitor in a cell-free biochemical assay^19^. Here, we placed this finding in a broader metabolic context by examining the dietary precursor of UA, EA, and the mixture of its major human phase II metabolites UA 3/8-gluc. In the ADP-Glo assay, UA inhibited DYRK1A with IC of 2.3 μM, in line with its reported ATP-competitive mechanism. In contrast to UA, its dietary precursor EA, did not influence DYRK1A activity at the tested concentration. Although EA is a known inhibitor of the structurally related kinase CK2, our results clearly show that its metabolic conversion to the urolithin dibenzo[*b*,*d*]pyran-6-ones scaffold is essential for emergence of DYRK1A inhibitory activity. Interestingly, glucuronidation, the major phase II metabolic modification of UA in humans, reduced its inhibitory potency by approximately one order of magnitude (IC_50_ ∼ 20 μM, but did not abolish activity. This shows that the UA tricyclic scaffold retains its ability to interact with the DYRK1A catalytic site after conjugation, albeit with reduced affinity.

The analysis of the previously published EA-CK2 X-ray data^42^ and the herein reported UA-DYRK1A cocrystal structure provides a molecular rationale for this SAR, particularly regarding the intriguing inability of EA to inhibit DYRK1A. The experimentally determined binding mode therefore provides a structural basis for understanding the activity of UA and its metabolites. Given the interest in urolithins and their derivatives as kinase modulators, the first experimentally determined crystal structure of UA bound to a protein target presented here provides a crucial reference for the structure-guided design of derivatives targeting DYRK1A and related kinases.

UA glucuronides partially retain the ability of the parent compound to inhibit of DYRK1A in the cell-free, enzymatic assay. However, a clear distinction between DYRK1A inhibition by UA and UA 3/8-gluc is seen at a cellular level. In a NanoBRET assay, UA demonstrated a robust TE, showing that it can access and engage DYRK1A under intracellular conditions. By contrast, although UA 3/8-gluc retained a measurable inhibitory activity in cell-free assay, no intracellular TE was seen. This difference is likely to reflect both the lower intrinsic affinity of the glucuronides for DYRK1A and their reduced cellular availability. Glucuronidation increases molecular polarity, enhances aqueous solubility, alters transporter-mediated disposition, generally facilitating elimination while limiting passive diffusion across cellular membranes^49^. On the other hand, numerous reports show *in vivo* effects of UA administration^8, 19^. Moreover, UA glucuronides may serve as a circulating reservoir of UA that can be locally deconjugated by β-glucuronidase (β-gluc), thereby recycling free UA. Deconjugation of UA glucuronides has been demonstrated under inflammatory conditions, supporting the possibility that biologically active UA may be released in specific tissue microenvironments^20^. Increased serum β-gluc has been previously reported in patients with DM^50^. However, whether this results in local deconjugation of UA glucuronides in pancreatic tissue remains unknown. Such mechanism could potentially contribute to the availability of free UA in DM-associated inflammatory conditions, but this possibility requires direct experimental validation.

Results of numerous *in vitro* and *in vivo* studies, as well as the few available human clinical trials, clearly suggest that UA may have a beneficial influence on human metabolism, including the potential to improve glucose homeostasis through previously unclear mechanisms. Notably, a prior screen of various polyphenols identified that both UA and UC enhance GSIS in INS-1 β-cells. However, the underlying pharmacological mechanism of activation of L-type Ca² channels^12^ and ERK1/2 pathway ^13^ was subsequently characterized specifically for UC, using isolated islets of Langerhans and perfused pancreas preparations, while the mechanisms responsible for the effect of UA were not further pursued. Here, we provide evidence supporting DYRK1A inhibition as a potential mechanism contributing to the effects of UA on β-cell function. In our cellular assays, UA increased both Ki67 proliferation and GSIS in MIN6 cells at low micromolar concentrations (1 and 2.5 µM), reaching levels comparable to those achieved with well-established, prototypic DYRK1A inhibitor harmine. Together with the biochemical, structural, and cellular TE data, these findings support a DYRK1A-mediated mechanism contributing to the effects of UA in MIN6 cells, consistent with those observed for harmine, and provide an additional mechanistic link distinct from the mechanisms characterized for UC. We show that UA promotes MIN6 cell proliferation, a phenotype not previously described for this compound, which is consistent with the established role of DYRK1A as a negative regulator of both β-cell replication and secretory function. This interpretation is consistent with a wealth of scientific literature in which the DYRK1A has been identified as a factor that supresses both β-cell proliferation and GSIS. Pharmacological DYRK1A inhibitors, including the prototypical inhibitor harmine, robustly promote human and rodent β-cell proliferation^14^. In addition, DYRK1A inhibition has been previously shown to enhance GSIS in MIN6 cells, the same model which was used in the present study.

Taken together, we show that UA and its glucuronides UA 3/8-gluc, but not its precursor EA, inhibit DYRK1A in a biochemical assay. We report the first crystal structure of UA bound to a protein target, providing a structural rationale for the observed SAR and a foundation for a rational design of derivatives with improved characteristics. Further, we show that UA inhibits DYRK1A in a cellular setting, and that this inhibition is associated with a concentration-dependent enhancement of β-cell proliferation and GSIS in MIN6 cells. These effects were comparable in magnitude to those of harmine, supporting DYRK1A inhibition as a mechanistic explanation for the previously described, but unexplained bioactivity of UA, consistent with the established role of the enzyme as a negative regulator of β-cell proliferation and function. Furthermore, the DYRK1A-mediated involvement of UA in the gut-microbe-DM/obesity axis may even be more multi-facetted, as recent findings suggest the involvement of DYKRs in increasing the number of incretin-expressing cells and improving glucose control in animal models^52^. Importantly, the first clinical trial has been initiated to evaluate whether prolonged UA administration enhances incretin secretion and improves glycemic control^53^. Therefore, the results presented here may contribute to a better understanding of the prospective outcomes of this trial.

Our findings provide a new perspective on the biological activity of UA, showing that DYRK1A is a molecular target involved in the regulation of pancreatic β-cell function by this compound. On the one hand, this mechanism may help to explain, at least in part, the beneficial effects attributed to wide variety of ellagitannin-rich foods in DM and related metabolic disorders, particularly in the context of pancreatic β-cell function. On the other hand, identification of DYRK1A as a putative target of UA may provide a rationale for the further development of pharmacological and nutritional strategies based on this metabolite or its derivatives. In this context, the newly developed, innovative synthetic route together with the presented, first insight into the structural features of UA interactions with its target protein, may facilitate future studies on SAR and support the design of urolithin-based compounds with improved biological properties and potential applications in the prevention or treatment of metabolic disorders.

## 4. Experimental Section

### 4.1. Synthesis of UA

Reagents and solvents were purchased from commercial suppliers and were used without additional purification. Thin layer chromatography (TLC) was performed using aluminum plates coated with silica gel (Silica gel 60 F254; grain size 0.040-0.063 mm, 230-400 mesh ASTM; Merck). After developing the plates in appropriate mobile phase systems (v/v), detection was performed using a UV lamp at the wavelength of 254 nm. Reverse-phase column chromatography (CC) separations were conducted using cartridge columns (12 g and 25 g SiliCycle SiliaSepTM C18). NMR spectra were recorded using the Agilent 400 MHz 400-MR DD2 self-service instrument. LC-MS analyses were performed using LC-MS 1260 Infinity II LC System coupled with the Agilent MSD 6100 mass detector: the Agilent 1260 Quaternary System chromatograph; the Agilent InfinityLab Poroshell 120 EC-C18 column, 3.0 x 50 mm, 2.7-Micron; thermostat temp. 35°C; UV detection at 220 and 250 nm and MS; Method: 0.5 min - 5% ACN/H2O + 0.1 % formic acid, 6.5 min-gradient to 95% ACN, + 0.1 % formic acid, 95% ACN + 0.1 % formic acid over 4 minutes. Fluorenone 3 is virtually insoluble in conventional organic solvents. Therefore, its samples for LC-MS analyses were dissolved in minimal amount of TFA and diluted in MeOH. From the same reason CF_3_CO_2_D was the solvent for NMR spectra registration of compound **3**. The observed proportion of product **3** to substrate **2** based on comparison of chromatographic peaks areas registered using UV detection at 220 nm were in accordance with those from the control ^1^H NMR analyses. The detailed synthetic protocols are provided in Supporting Information.

### 4.2. Biochemical and biophysical evaluation of DYRK1A inhibition

#### 4.2.1. ADP-Glo kinase activity assay

The inhibitory potency (IC_50_) of harmine, EA, UA and UA 3/8-gluc^36^ against DYRK1A kinase domain was determined using the ADP-Glo kinase activity assay (Promega). The synthetic peptide DYRKtide (RRRFRPASPLRGPPK, Caslo ApS) was used as a substrate for DYRK1A. Briefly, 20 µL of kinase in reaction buffer (100 mM MOPS, pH 6.8, 100 mM KCl, 10 mM MgCl_2_) was incubated with 10 µL of compound for 15 minutes at room temperature. The reaction was initiated by the addition of 20 µL of substrates mixture (ATP and DYRKtide) and allowed to proceed for 30 minutes at room temperature. Subsequently, 5 µL of reaction mixture was transferred to HTRF 96-well low-volume plates (Cisbio), and ATP consumption was measured with ADP-Glo Kinase Assay kit (Promega) according to manufacturer’s instructions. The final assay conditions were as follows: DYRK1A at 5 ng/µl, compound (EA, UA, UA3/8-gluc or harmine) at 20 μM for single-dose inhibition or in concentration range of 40 nM –20 µM for IC_50_ determination, ATP at 100 µM and DYRKtide at 400 µM. All measurements were performed in triplicate, and IC_50_ values were determined using GraphPad Prism software.

#### 4.2.2. Nano-Differential Scanning Fluorimetry (nanoDSF)

DYRK1A kinase domain stability in the presence of harmine, EA, UA, and UA 3/8-gluc was assessed by determining protein melting temperatures determination using nano-differential scanning fluorimetry (nanoDSF) on Prometheus NT.48 instrument (NanoTemper Technologies). Briefly, DYRK1A protein (1.5 mg/ml) was incubated with each compound (10 μM) or DMSO as a control in 100 mM MOPS, pH 6.8, 100 mM KCl, 10 mM MgCl_2_ for 15 minutes at room temperature. Samples were loaded into high-sensitivity premium capillaries and subjected to temperature ramp from 20°C to 95°C at a rate 1°C/min, while intrinsic tryptophan fluorescence at 330 and 350 nm was recorded. The melting temperature (Tm) was determined as the inflection point of the fluorescence ration (F350/F330) as a function of temperature. All experiments were performed in triplicates.

#### 4.2.3. Nano Bioluminescence Resonance Energy Transfer (BRET) Target Engagement (TE) Assay

The inhibitory potency (IC_50_) of harmine, UA and UA 3/8-gluc against DYRK1A kinase in complex cellular context was determined using NanoBRET^TM^ Target Engagement Intracellular Kinase Assay, K-10 (Promega), according to the manufacturer’s instructions. Briefly, HEK293 cells (American Type Culture Collection, ATCC CRL-1573) were transiently transfected with NanoLuc-DYRK1A Fusion vector (Promega) using FuGENE^®^ HD transfection reagents (Promega). The vector encodes NanoLuc luciferase fused to the N-terminus of full-length human DYRK1A. 48 hours post-transfection, cells were incubated with the NanoBRETTM K-10 tracer (0.5 μM final concentration) in Opti-MEM medium for 15 minutes at 37°C, followed by the addition of tested compounds (15 nM - 200 μM final concentration range) and further incubation for 2 hours at 37°C. Subsequently, NanoBRET™ NanoGlo^®^ substrate and extracellular Nanoluc were added in Opti-MEM according to the manufacturer’s protocol, and BRET signals were measured using GloMax^®^ Discovery Plate Reader (Promega). All measurements were performed in triplicate, and IC_50_ values were determined using GraphPad Prism software.

### 4.3. Protein crystallization, data collection, and structure determination

Recombinant DYRK1A kinase domain was concentrated to 10–13 mg·mL ¹ and incubated for 1 h at 20°C with a fivefold molar excess of UA under gentle agitation (600 rpm). For crystallization screening, the protein–ligand complex was mixed with reservoir solutions in a 0.45:0.30 (v/v) ratio. Crystallization trials were performed at 20°C using the sitting-drop vapor diffusion method. Initial crystals appeared within 1–5 days. Diffraction-quality crystals were obtained under Morpheus B2 conditions (Molecular Dimensions), consisting of a precipitant mix of polyethylene glycols, salts, and buffer system as provided by the manufacturer. Crystals were cryoprotected by transferring them into mother liquor supplemented with 25% (v/v) ethylene glycol and subsequently flash-cooled in liquid nitrogen. X-ray diffraction data were collected at the BESSY II synchrotron (Berlin, Germany). Diffraction data were processed using XDS^54^ for indexing and integration, followed by scaling with AIMLESS^55^ within the CCP4i2 suite.^56^ Structure solution was carried out by molecular replacement using PHASER^57^ with the DYRK1A structure (PDB ID: 6EIS) as the search model. Subsequent model building and refinement were performed using PHENIX,^58^ including iterative cycles of phenix.refine^59^ and manual building in Coot.^60^ Restraints for urolithin A were generated using the GRADE Web Server13^61^ and incorporated during refinement. The final model was validated using standard crystallographic validation tools implemented in PHENIX.

### 4.4. Cellular studies

#### 4.4.1. Materials for cell culture

Materials for cell culture were obtained from commercial suppliers: DMEM high glucose (4.5 g/L) (#11995-065, Gibco), DMEM low glucose (1.0 g/L) (#P04-01550, PAN Biotech), FBS (#A5670801, Gibco), penicillin-streptomycin (10,000 U/mL) (#15140122, Gibco), β-mercaptoethanol (βME) 100× (#ES-007-E, Sigma-Aldrich), ciprofloxacin (#17850-5G-F, Sigma-Aldrich), DPBS (#14040117, Gibco), TrypLE Express (#12604021, Gibco).

#### 4.4.2. Cytotoxicity study

MIN6 β-cells were seeded in a 96-well plate at a density of 1×10^4^ cells/well in a complete medium consisting of high-glucose DMEM (4.5 g/L), supplemented with 15% FBS, 1% penicillin-streptomycin (P/S), 55 µM β-mercaptoethanol (β-ME), and 10 µg/mL ciprofloxacin. After 24 h of seeding, cells were treated with harmine (2.5 µM) and different concentrations of UA (1, 2.5, 10, and 25 µM) and incubated for 24 h. The medium was subsequently replaced with fresh medium, and cells were further incubated for 48 h. At the end of treatments, the medium was replaced with the phenol red-free DMEM containing Alamar Blue (final concentration of 44 µM) to each well. Then the plate was co-incubated for 3 to 4 h to allow color development, following fluorescence measurement with the Promega GloMax® Discover microplate reader at excitation of 520 nm and emission of 580-640 nm. The percentage of cell viability was normalized to control (untreated) as 100% viable cells. A viability threshold (<80% relative to control) was used to indicate significant cytotoxic effects.

#### 4.4.3. Glucose-stimulated insulin secretion (GSIS) and Enzyme-Linked Immunosorbent Assay (ELISA) experiments

MIN6 β-cells were seeded in a 96-well plate at a density of 1×10^4^ cells/well in the complete medium. After 24 h of seeding, cells were treated with harmine (2.5 µM) or UA (1, 2.5 µM) and incubated for 24 h. The medium was subsequently replaced with fresh medium, and cells were further incubated for 24 h. Then, GSIS assay was performed to assess insulin secretion. The medium was first replaced with complete low-glucose medium (DMEM 1.0 g/L, ∼5.5 mM) and the cells were incubated for 24 h. At the end of this pre-incubation, cells were first incubated with Krebs-Ringer bicarbonate (KRB) buffer containing very low glucose (2 mM) for 2 h, followed by incubation to high-glucose KRB (25 mM) for 30 min, with supernatants collected. At the end of the procedure, cells were lysed with the RIPA buffer, and the lysates were also collected. Insulin concentrations were quantified by using the Invitrogen Insulin Mouse ELISA Kit (#EMINS, Thermo Fisher Scientific) according to the manufacturer’s protocol, and the absorbance was read out with the Promega GloMax® Discover microplate reader at 450 nm. Insulin concentrations were calculated by a four-parameter logistic curve, and results were normalized to control.

#### 4.4.4. Cell proliferation assay

MIN6 β-cells were seeded in a 96-well plate at a density of 1×10^4^ cells/well. After 24 h of seeding, cells were treated with harmine (2.5 µM) and low concentrations of UA (1 and 2.5 µM) and incubated for 24 h. The medium was subsequently replaced with fresh medium, and cells were further incubated for 72 h. Cells were washed with PBS and then fixed using 4% paraformaldehyde for 10-15 min at room temperature to preserve cellular structures. Fixed cells were permeabilized with 0.15% Triton X-100 for 10 min to allow intracellular antibody access. Cells were incubated in blocking buffer (1% BSA in PBS) for 30 min to minimize non-specific antibody binding. Cells were incubated for 1.5 h at 37°C with Ki-67 Monoclonal Antibody (SolA15), eFluor™ 570 eBioscience™ (#41-5698-82, Invitrogen) and Insulin Monoclonal Antibody, Alexa Fluor™ 488, (#53976982, eBioscience™), both diluted to 100× in the blocking buffer. Cells were washed and incubated with DAPI (0.5 µg/mL) for 10 min and washed five times with PBS. Cells were imaged and visualized with a Zeiss Celldiscoverer 7 (Zeiss, Jena, Germany) and captured images were processed using imageJ software.

## Supporting information

Synthetic details, Supplemantary Figures S1, S2 and Supplementary Table 1

## Author contributions

M. N.: conceptualization, investigation, methodology, writing – original draft, writing – review and editing; A. B.: investigation, data curation, writing – original draft, writing – review and editing, visualization; K. P.: conceptualization, investigation, data curation, methodology, validation, writing – original draft, writing – review and editing, visualization; A. Mar: investigation, data curation, writing – original draft, writing – review and editing, visualization; M. P. R.: Investigation, data curation, writing – review and editing; A. Mat: investigation, data curation, writing – review and editing; P. G.: investigation, data curation, methodology; supervision; writing – review and editing; J. P.: resources, writing – original draft, writing – review and editing; A. C.: conceptualization, funding, methodology, resources, supervision, visualization, writing – original draft, writing – review and editing; M. D.: conceptualization, data curation, funding, methodology, project administration, resources, supervision, writing – original draft, writing – review and editing, visualization.

## Acknowledgements

The Authors would like to thank Dr Tomasz Kantyka (Malopolska Centre of Biotechnology, Jagiellonian University, Krakow, Poland) for providing access to the Zeiss Celldiscoverer 7 and Ms. Beril Anita Yiğit for her assistance with preliminary experiments. The Structural Biology Core Facility of Małopolska Centre of Biotechnology is acknowledged for valuable support. We thank Helmholtz-Zentrum Berlin for the allocation of neutron/synchrotron radiation beamtime (BESSY Beamline 14.1). This work was supported by grants from the National Science Centre (UMO-2022/45/B/NZ7/04269) to A. C. and (UMO-2025/57/N/NZ1/03364) to A. B., respectively.

## Conflicts of Interest

The authors declare no conflicts of interest.

## Data Availability Statement

The data that support the findings of this study are available from the corresponding authors upon reasonable request.

## Supporting Information

Supporting Information is available from the publisher or from the author.

## Notes

### Competing Interest Statement

The authors have declared no competing interest.

