## Supplementary material for "DYRK1A inhibition by the postbiotic metabolite Urolithin A contributes to enhanced β-cell function: Structural, Biochemical and Cellular Evidence": Synthetic details, Supplemantary Figures S1, S2 and Supplementary Table 1

**1. Synthetic protocols**

**1.1. Synthesis of 2,7-diacetylfluorene 2**

The compound was obtained by modifying the literature procedure.^1^

An oven-dried, round-bottom bulb (500 mL) was charged with AlCl_3_ (64.18 g, 481.3 mmol, 8 equiv) and anhydrous DCM (120 mL). The stirred suspension was cooled down to -10°C and Ac_2_O (22.8 mL, 240.6 mmol, 4 equiv) was added dropwise. A solution of fluorene **1** (10.00 g, 60.16 mmol, 1 equiv) in of anhydrous DCM (60 mL) was added dropwise to the resulting suspension and the cooling bath was removed. The reaction mixture was stirred at rt for 3 h, cooled (NaCl_aq_-CO_2_ bath) and poured slowly to cooled 1M HCl_aq_ (300 mL). The layers were separated and the aqueous phase was extracted with DCM (3 x 160 mL). The combined organic extract were washed with brine (160 mL). Brine was re-extracted with DCM (100 mL) and the combined organic extract was washed with saturated aqueous NaHCO_3_ until evolution of CO_2_ ceased and the pH of aqueous phase reached ~8. The layers were separated and the aqueous one was re-extracted with DCM (75 mL). All organic extracts were combined and dried over anhydrous Na_2_SO_4_, filtrated and evaporated to obtain 15.077 g (~100%) of the crude product, the purity (> 95%) of which was sufficient for further experiments.

^1^H NMR (400 MHz, CDCl_3_) δ 8.17 (s, 2H), 8.03 (d, *J* = 8.0 Hz, 2H), 7.90 (d, *J* = 8.0 Hz, 2H), 4.01 (s, 2H, CH_2_), 2.67 (s, 6H, 2 x CH_3_). LR MS *m/z*  251 [M + H]^+^, 523 [2M + Na]^+^.

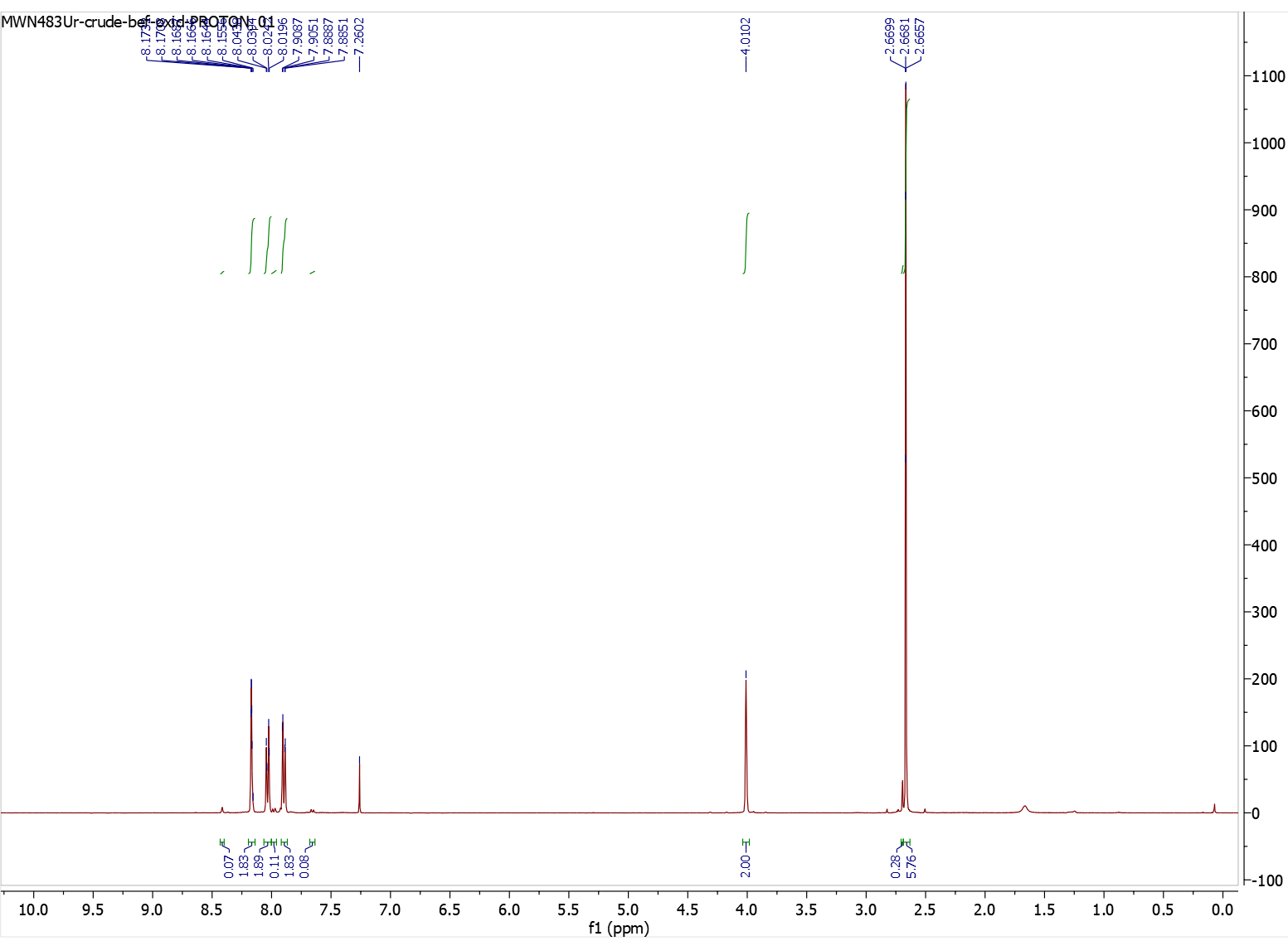

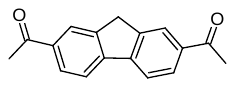

**1.2. Synthesis of 2,7-diacetyl-9*H*-fluoren-9-one 3 by the K_2_Cr_2_O_7_-mediated oxidation of 2.**

The compound was obtained by modifying the literature procedure.^2^

A round-bottom flask (250 mL) was charged with 2,7-diacetylfluorene **2** (5.500 g, 22.0 mmol, 1 equiv.), AcOH (110 mL) and K_2_Cr_2_O_7_ (8.598 g, 29.23 mmol, 1 .33 equiv.). The resulting mixture was heated at reflux for 3.5 h, cooled down to rt, and the precipitate was filtered off and washed with AcOH (5 times) and water (3 times). The precipitate was suspended in a MeOH/DCM mixture (200 mL, 1:1, v/v) and DCM was evaporated using a rotary evaporator. The resulting solid was filtered and washed with MeOH to give 5.11 g (88%) of the title compound as a light green solid.

^1^H NMR (400 MHz, CF_3_CO_2_D) δ 8.35 (s, 2H, C^1^H & C^8^H) overlapping 8.29 (d, *J* = 7.2 Hz, 2H, C^3^H & C^6^H), 7.77 (d, *J* = 7.8 Hz, 1H, C^4^H & C^5^H), 2.71 (s, 6H, 2 x CH_3_); ^13^C NMR (101 MHz, CF_3_CO_2_D) δ 205.8 (2 x C=O), 197.8 (2 x C=O), 150.9, 140.2, 139.5, 136.9, 127.7, 124.2, 27.0 (2 x CH_3_). LR MS m/z 265 [M + H]^+^, 552 [2M + Na]^+^.

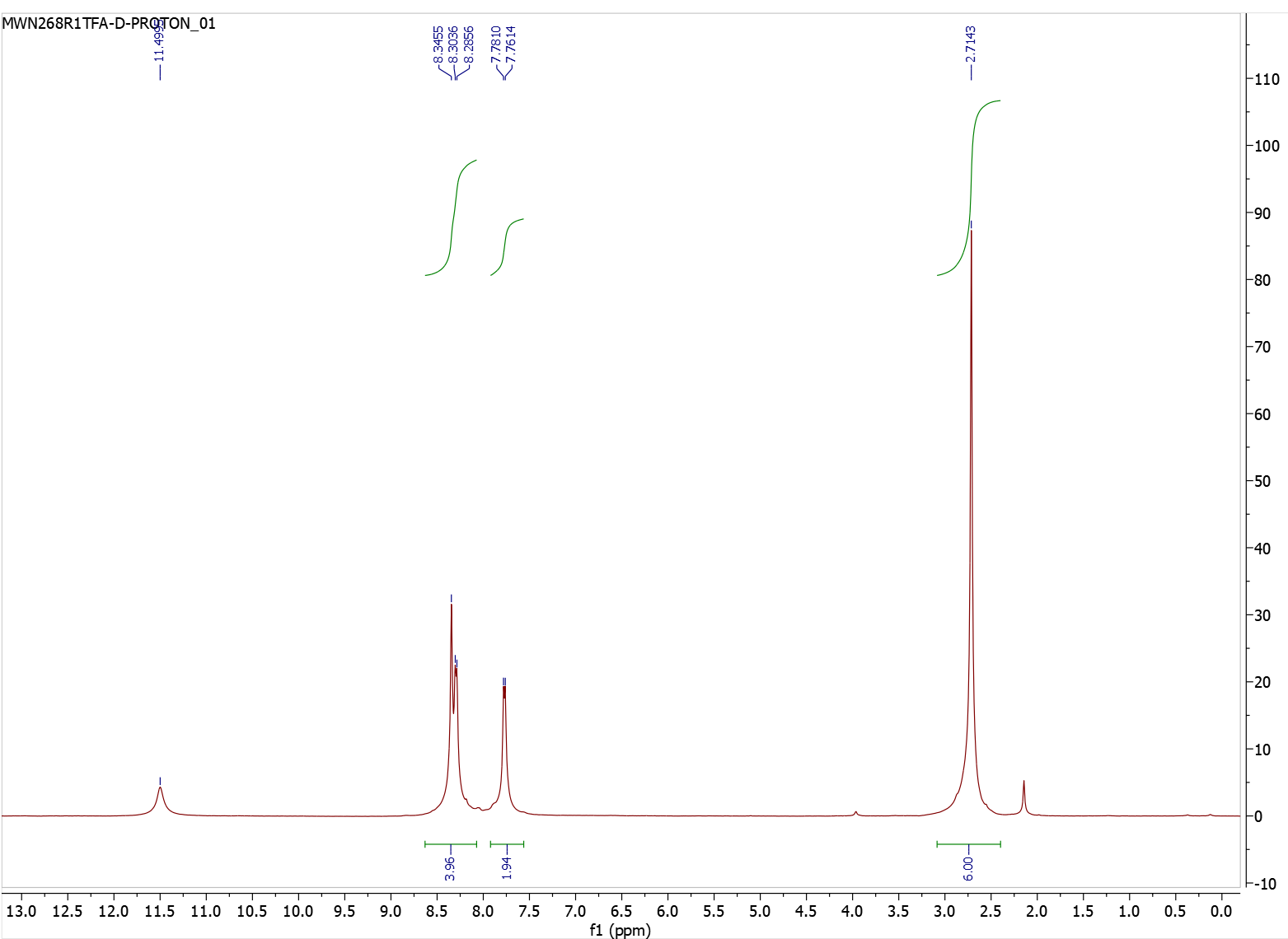

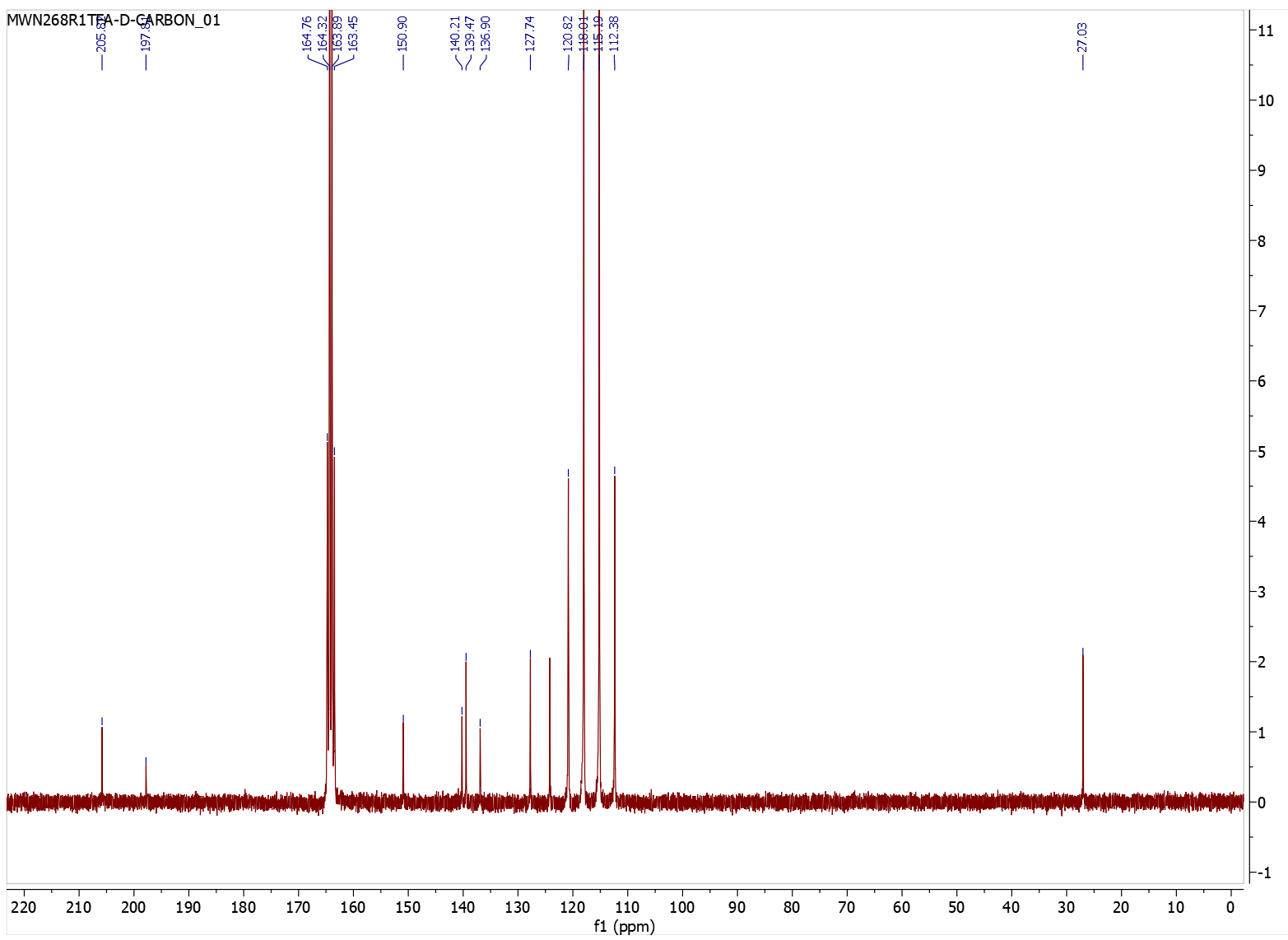

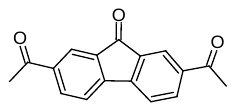

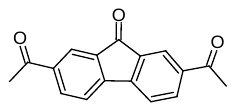

**1.3. Synthesis of 2,7-diacetyl-9*H*-fluoren-9-one 3 by oxidation of 2 with molecular oxygen involving the SET mechanism.**

A round-bottom flask (250 mL) was charged with 2,7-diacetylofluorene **2** (3.000 g, ~12 mmol, 1 equiv), and DMF (120 mL). The suspension was stirred at rt until dissolution and then placed in the heating bath (30°C). K_2_CO_3_ (166 mg, 1.2 mmol, 10 mol%) was added and the mixture was stirred at 30°C for 4 days in open flask. The mixture was cooled down to rt and placed in a refrigerator (5°C) overnight. The precipitated solid was filtrated, washed with chilled DMF (3 x 4 mL), water (5 x 5 mL) and dried under high vacuum to obtain 2.455 g of crude product (product/substrate ratio: 92:8; LC-MS-UV at 220 nm). This material was placed in a pressure vessel, DCM (100 mL) was added and the resulting suspension was refluxed for 75 min at 60°C, cooled down to rt and kept in the fridge overnight. The solid was filtrated, washed with cooled DCM (3 x 5 mL) and dried *in vacuo* to obtain 1.925 g (61% from 2 steps) of product **3** as an orange solid (purity LC-MS UV 220 nm; 99%).

Additional crop of product was obtained as follows: The DMF filtrate from the first crop was mixed with water (107 mL). The resulting suspension was kept in the refrigerator (5°C) overnight, the solid was collected using Schott funnel, washed with DMF (3 x 5 mL) and water (4 x 5 mL), and air-dried. The solid was combined with the residue obtained by the evaporation of DCM filtrate from the first crop. This material was placed in a pressure vessel, suspended in 50 mL of DCM, heated for 2 h at 60°C, cooled to rt and kept in the refrigerator (5°C) overnight. The resulting solid was filtrated and washed with chilled DCM (4 x 2 mL) to give additional 355 mg of product (purity LC-MS UV 220 nm; 96%) as an orange solid. Overall yield: 2.228 g (72% from 2 steps).

**1.4. Synthesis of 3,8-dihydroxy-6*H*-benzo[*c*]chromen-6-one UA, 0.57 mmol scale.**

A glass vial (24 mL) with a Teflon-sealed screw-cap was charged with 2,7-diacetyl-9*H*-fluoren-9-one **3** (150 mg, 0.57 mmol, 1 equiv) and CF_3_CO_2_H (1.5 mL, 35 equiv). The resulting solution was cooled to -15°C, followed by dropwise addition of 30% aqueous solution of H_2_O_2_ (0.87 mL, 15 equiv). After 10 min, the reaction vial was placed in refrigerator (5°C) for 64 h. The reaction mixture was cooled to 0°C and cold, distilled water (5 ml) was added dropwise. The reaction mixture was added to a cooled suspension of NaHCO_3_ (1.907 g, 22.7 mmol, 40 equiv) in distilled water (5 ml). The suspension was extracted with AcOEt (4 x 15 ml). The organic extracts were combined, washed with brine (10 ml), dried over Na_2_SO_4_, filtered and concentrated using a rotary evaporator. MeOH (1 mL) and 1 M aqueous HCl (1 mL) were added to the residue and the resulting mixture was stirred at 70°C for 4 h. The volatiles were evaporated, and the residue was triturated with AcOEt (1.5 mL), filtered, and washed sequentially with three small portions of cold AcOEt and *n*-hexane to give 51 mg (39%) of the title product as a white solid.

^1^H NMR (400 MHz, DMSO-*d*_6_) δ 10.19 (s, 1H, OH), 10.13 (s, 1H, OH’), 8.11 (d, *J* = 8.8 Hz, 1H), 8.02 (d, *J* = 8.7 Hz, 1H), 7.51 (d, *J* = 2.7 Hz, 1H), 7.32 (dd, *J* = 8.7, 2.7 Hz, 1H), 6.80 (dd, *J* = 8.7, 2.4 Hz, 1H), 6.72 (d, *J* = 2.4 Hz, 1H). LR MS *m/z*  229 [M + H]^+^, 479 [2M + Na]^+^.

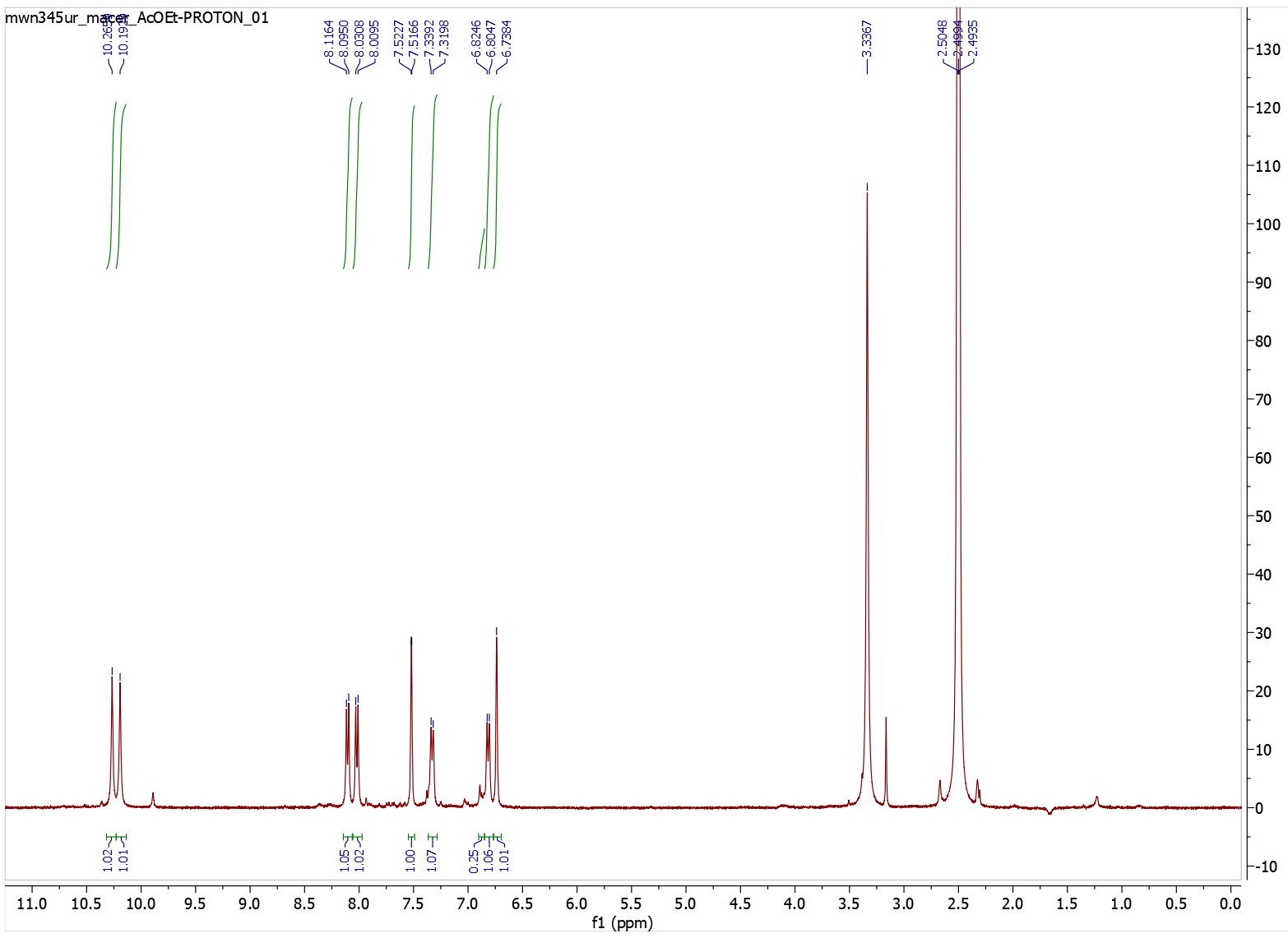

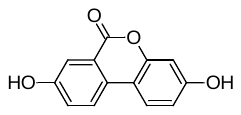

**1.5. Synthesis of 3,8-dihydroxy-6*H*-benzo[*c*]chromen-6-one UA, 4.00 mmol scale.**

A screw-cap vial (60 mL) was charged with 2,7-diacetyl-9*H*-fluoren-9-one **3** (1.057 g, 4.00 mmol, 1 equiv) and TFA (10.7 mL) was added. The mixture was stirred at rt until dissolution of the solids and cooled down to -15°C. To the stirred, cooled solution was added dropwise 30% aqueous solution of H_2_O_2_ (6.1 mL, 60 mmol H_2_O_2_, 15 equiv), over 5 min, and the reaction mixture was stirred at 5°C (*i*-PrOH; immersion cooler) for 70 h. The resulting suspension was cooled down to 0°C and cold, distilled water (35 ml) was added dropwise during 15 min. The mixture was stirred at rt for 23 h, which resulted in a partial hydrolysis of acetyl groups. The mixture was cooled (0°C) and added dropwise to a stirred, cooled (0°C) suspension of NaHCO_3_ (13.440 g 160 mmol, 40 equiv) in distilled water (35 mL) and PhMe (50 mL). Further PhMe (50 mL) and AcOEt (50 mL) were added and the layers were separated using separatory funnel. The aqueous phase was extracted with AcOEt (7 x 50 mL). The organic extracts were combined, washed with brine (75 mL) dried over Na_2_SO_4_, filtrated and evaporated. The crude product was refluxed with THF (30 mL) until complete dissolution, followed by slow evaporation until 1/3 of primary volume (~10 mL) using rotary evaporator (without immersing in the heating bath) until the precipitation of solid was observed. The mixture was filtrated, washed with chilled THF (4 x 0.5 mL), *n*-pentane (2 x 1 mL), and dried in vacuo to obtain 200 mg (22%) of product as a beige solid.

Additional crop of product was obtained as follows: The THF filtrate containing significant amount of incompletely hydrolysed **UA** was concentrated *in vacuo*. To the residue was added THF (10 mL), 1M aqueous HCl (10 mL) and the mixture was heated at 70°C for 3 h. The solution was flushed with argon and concentrated using rotary evaporator. To the residue was added AcMe (10 mL) and the suspension was stirred at 60°C for 1 h and left overnight to cool to rt. The resulting suspension was filtrated, washed with chilled AcMe (4 x 0.5 mL) to obtain additional 105 mg of product as a beige solid.

The product can be further recovered as follows: The filtrate was evaporated and subjected to reverse phase column chromatography (C-18; using THF/H_2_O 25-35%). The purest fractions were collected, concentrated, dissolved in THF and filtrated through a plug of cotton and concentrated *in vacuo*. To the crude was added AcMe (5 mL) and the mixture was heated at 60°C for 1 h, left over night to cool down to rt, filtrated, washed with chilled AcMe (4 x 0.5 mL) and dried in vacuo to obtain additional 36 mg of a product as a beige solid. Overall yield 341 mg (37%).

^1^H NMR (400 MHz, DMSO-*d*_6_) δ 10.16 (br s, 2H, 2 x OH), 8.09 (d, *J* = 8.8 Hz, 1H), 8.00 (d, *J* = 8.7 Hz, 1H), 7.51 (d, *J* = 2.7 Hz, 1H), 7.31 (dd, *J* = 8.7, 2.7 Hz, 1H), 6.80 (dd, *J* = 8.7, 1.6 Hz, 1H), 6.72 (d, *J* = 1.8 Hz, 1H); ^13^C NMR (101 MHz, DMSO-*d*_6_) δ 160.6, 158.6, 157.0, 150.9, 127.0, 124.1, 123.8, 123.6, 120.2, 113.6, 113.0, 109.9, 102.9. LR MS *m/z*  229 [M + H]^+^, 479 [2M + Na]^+^.

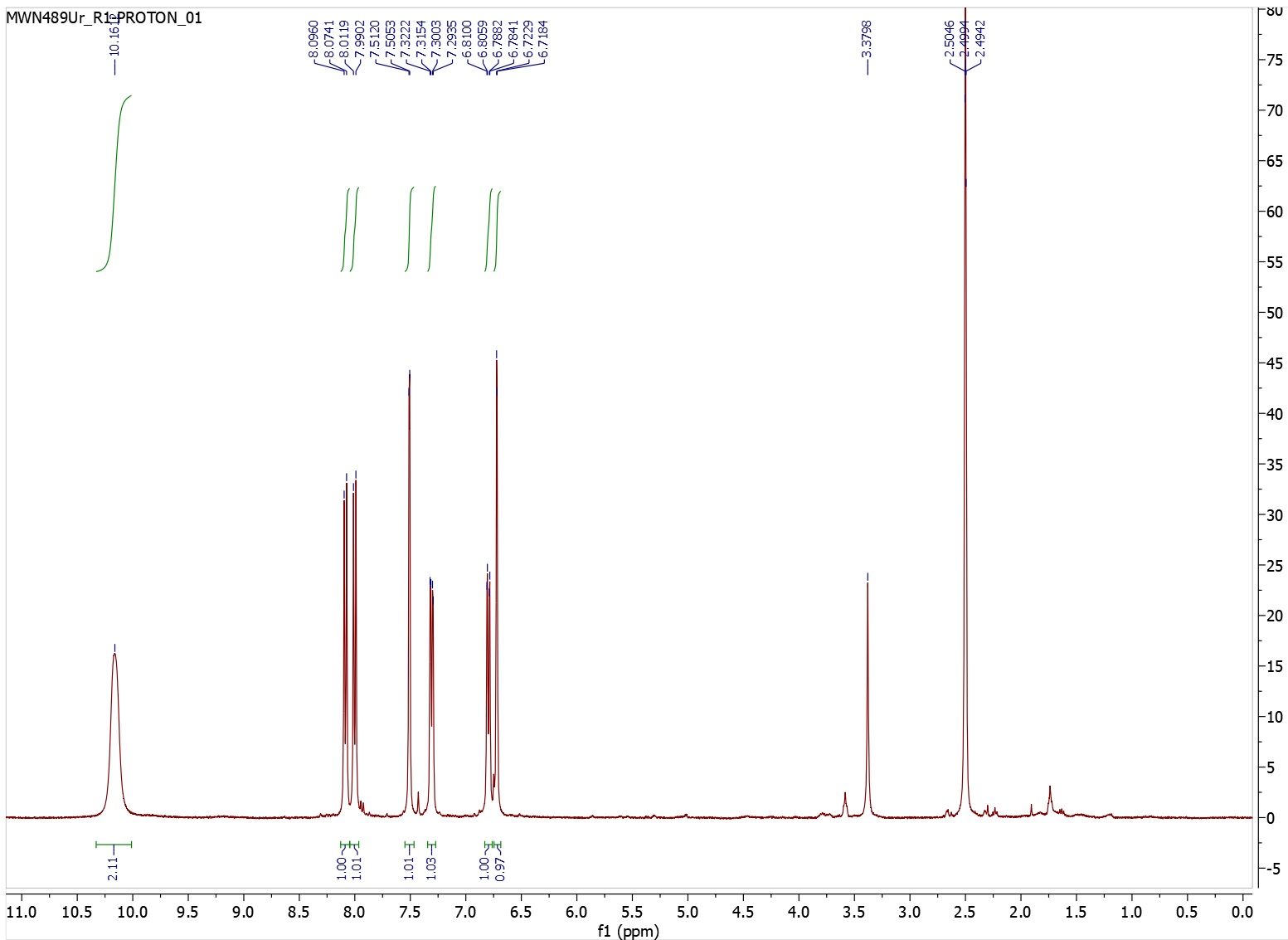

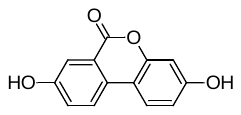

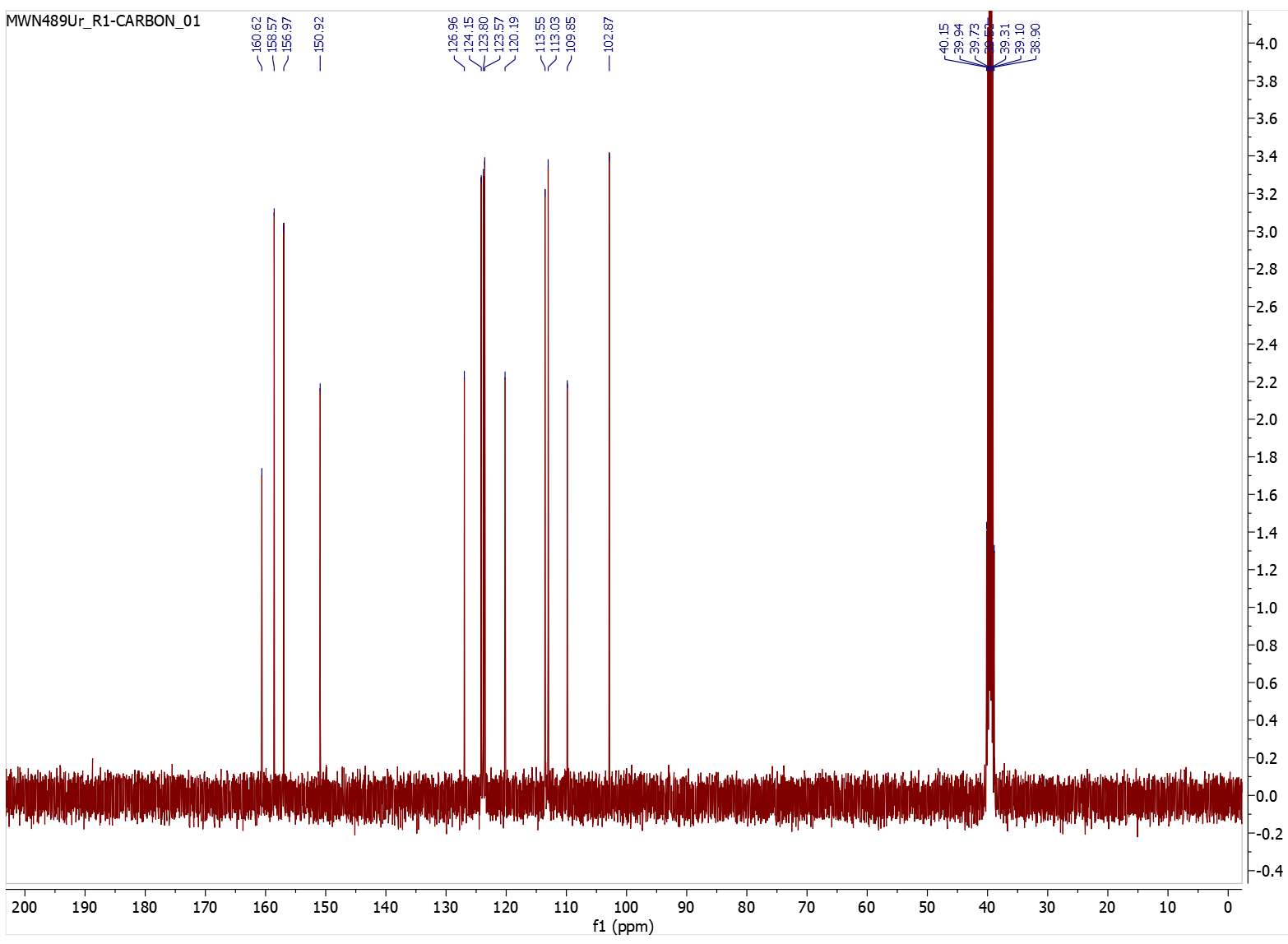

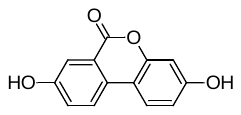

**1.6. Attempt to synthesize UA from 2, by employing m-CPBA as a Baeyer-Villiger oxidant. Synthesis of 2,7-diacetoxy-9*H*-fluoren-9-one**

A screw-capped vial (60 mL) was charged with 2,7-diacetylfluorenone **2** (528.5 mg, 2.00 mmol, 1 equiv) and CHCl_3_ (20 mL). The suspension was stirred at rt, ~77% *m*-CPBA stabilized with water (2.689 g, 12 mmol, 6 equiv) was added portionwise and the reaction mixture was stirred at rt for 5 days in closed vial protected from light using aluminum foil. Saturated aqueous solution of NaHCO_3_ 20 mL) was added dropwise and the mixture was stirred at rt for 1h. 2 M aqueous solution of Na_2_S_2_O_3_ (6 mL) was added and the mixture was stirred at rt for additional 1 h. The phases were separated and the aqueous one was washed with CHCl_3_ (4 x 20 mL). The organic extracts were combined washed with saturated aqueous solution of NaHCO_3_ (5 mL) and brine (20 mL). The brine was reextracted with CHCl_3_ (20 mL). All organic extracts were combined, dried over Na_2_SO_4_ and evaporated. The crude product was dissolved in hot DCM and subjected to column chromatography (silica; using DCM 100%) to give 410 mg (69%) of product as a yellow solid.

^1^H NMR (400 MHz, DMSO-*d*_6_) δ 7.84 (dd, *J* = 8.0, 0.3 Hz, 2H), 7.42-7.39 (m, 2H), 7.37 (dd, *J* = 8.0, 2.2 Hz, 2H), 2.28 (s, 6H, 2 x CH_3_). The spectrum is in accordance with literature.^3^; ^1^H NMR (400 MHz, CDCl_3_) δ 7.49 (d, *J* = 8.0 Hz, 2H), 7.38 (d, *J* = 2.1 Hz, 2H), 7.20 (dd, *J* = 8.0, 2.1 Hz, 2H), 2.32 (s, 6H, 2 x CH_3_); ^13^C NMR (101 MHz, CDCl_3_) δ 191.6 (C=O), 169.3, 151.5, 141.4, 136.0, 127.8, 121.2, 118.5, 21.2 (2 x CH_3_). LR MS *m/z*  297 [M + H]^+^, 593 [2M + Na]^+^. 615 [2M + Na]^+^.

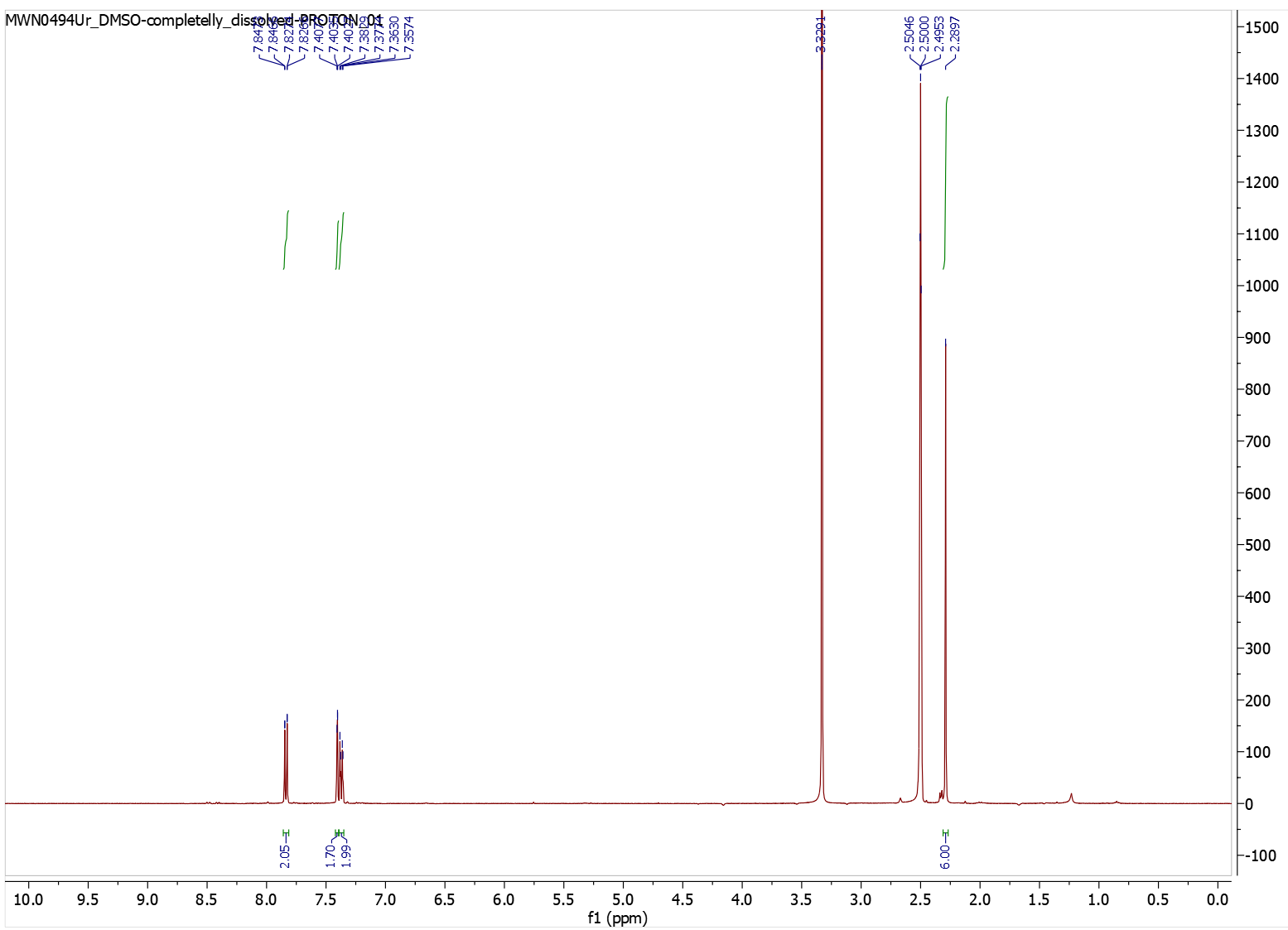

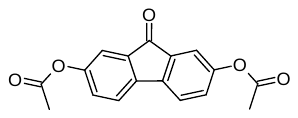

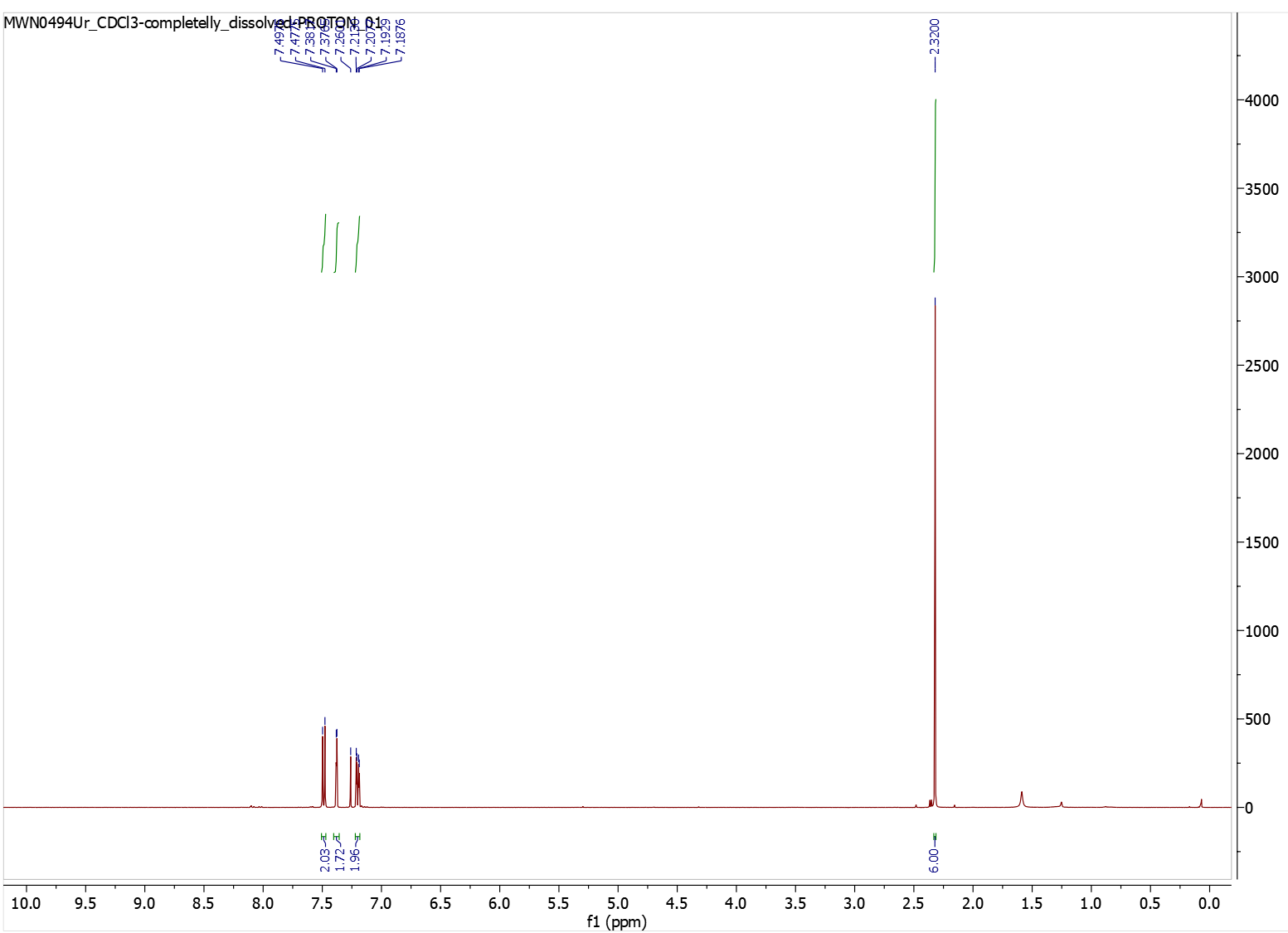

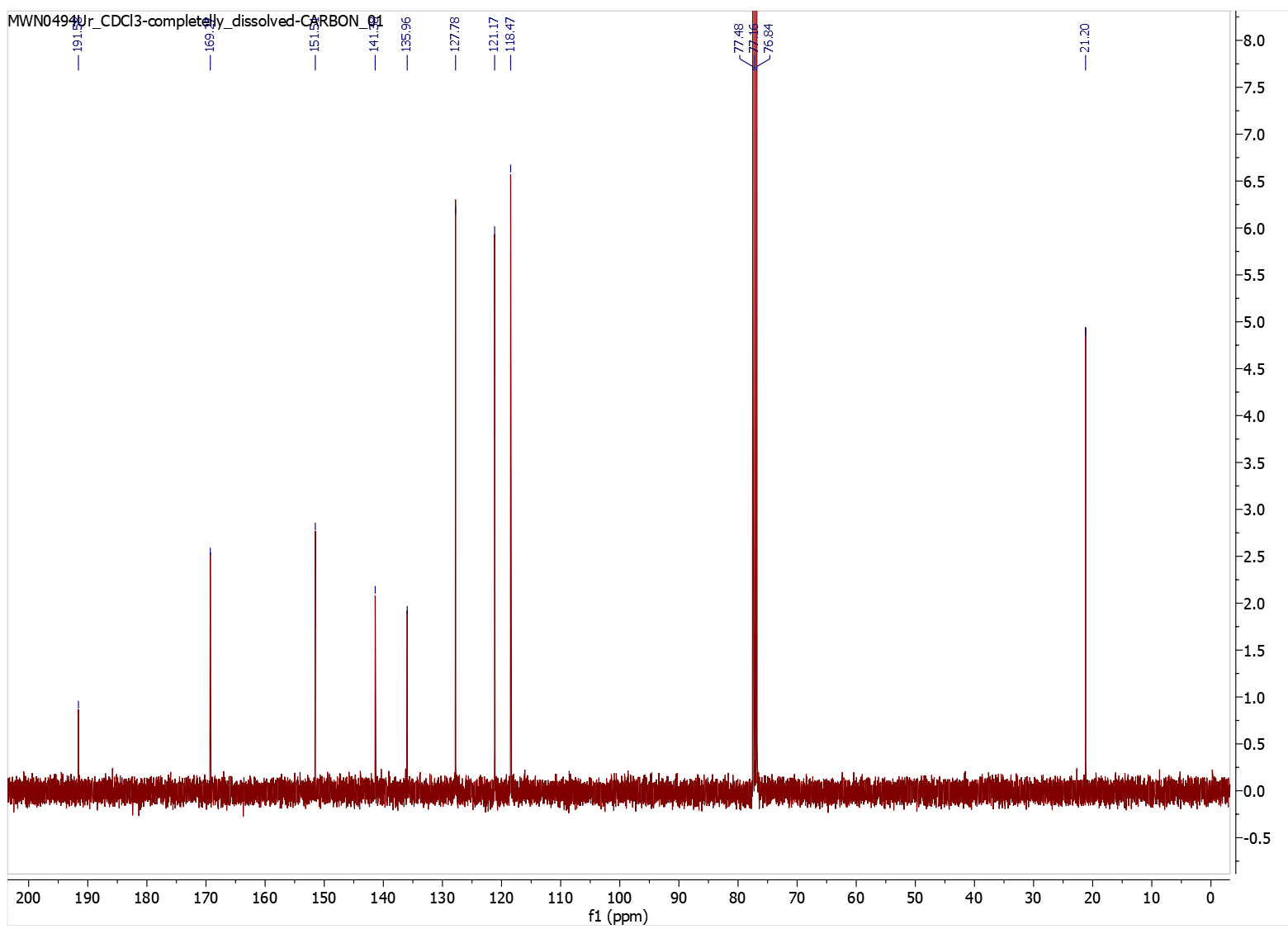

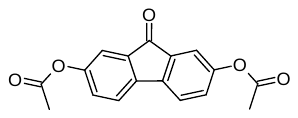

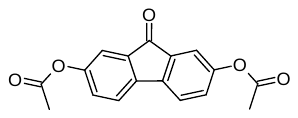

**1.7. Attempt to synthesize UA from directly from 2,7-dihydroxy-9*H*-fluoren-9-one**

A screw-cap vial (8 mL) was charged with 2,7-dihydroxy-9*H*-fluoren-9-one (30 mg, 0.14 mmol, 1 equiv) and TFA (0.38 mL, 4.95 mmol) was added. The suspension was stirred at rt for 10 min. and cooled to -15°C. 30% aqueous solution of H_2_O_2_ (0.22 mL, 2.12 mmol H_2_O_2_, 15 equiv) was added dropwise, over 1 min, and the reaction mixture was stirred at +5°C for 70 h. The resulting suspension was cooled down to 0°C and cold, distilled water (1 ml) was added dropwise. The resulting suspension was added dropwise to a screw-cap vial (24 mL) containing a suspension of NaHCO_3_ (475 mg, 5,66 mmol, 40 equiv) in distilled water (1 mL), while maintaining the temperature ~0°C. To the suspension was added AcOEt (4 mL) and, after 10 min. of stirring, the phases were separated. The aqueous layer was extracted with AcOEt (3 x 4 mL). All organic extracts were combined, washed with brine (2 mL) and the phases were separated. The organic phase was dried over Na_2_SO_4_, filtrated, evaporated and dried *in vacuo* to obtain crude, brown solid. ^1^H NMR analysis of this material showed only the unreacted substrate.

^1^H NMR (400 MHz, DMSO-*d*_6_) δ 9.86 (s, 2H, 2 x OH), 7.37 (d, *J* = 7.9 Hz, 2H), 6.88 (d, *J* = 2.0 Hz, 2H), 6.86 (dd, *J* = 7.9, 2.3 Hz, 2H). 447 [2M + Na]^+^. The spectrum is in agreement with literature.^4^

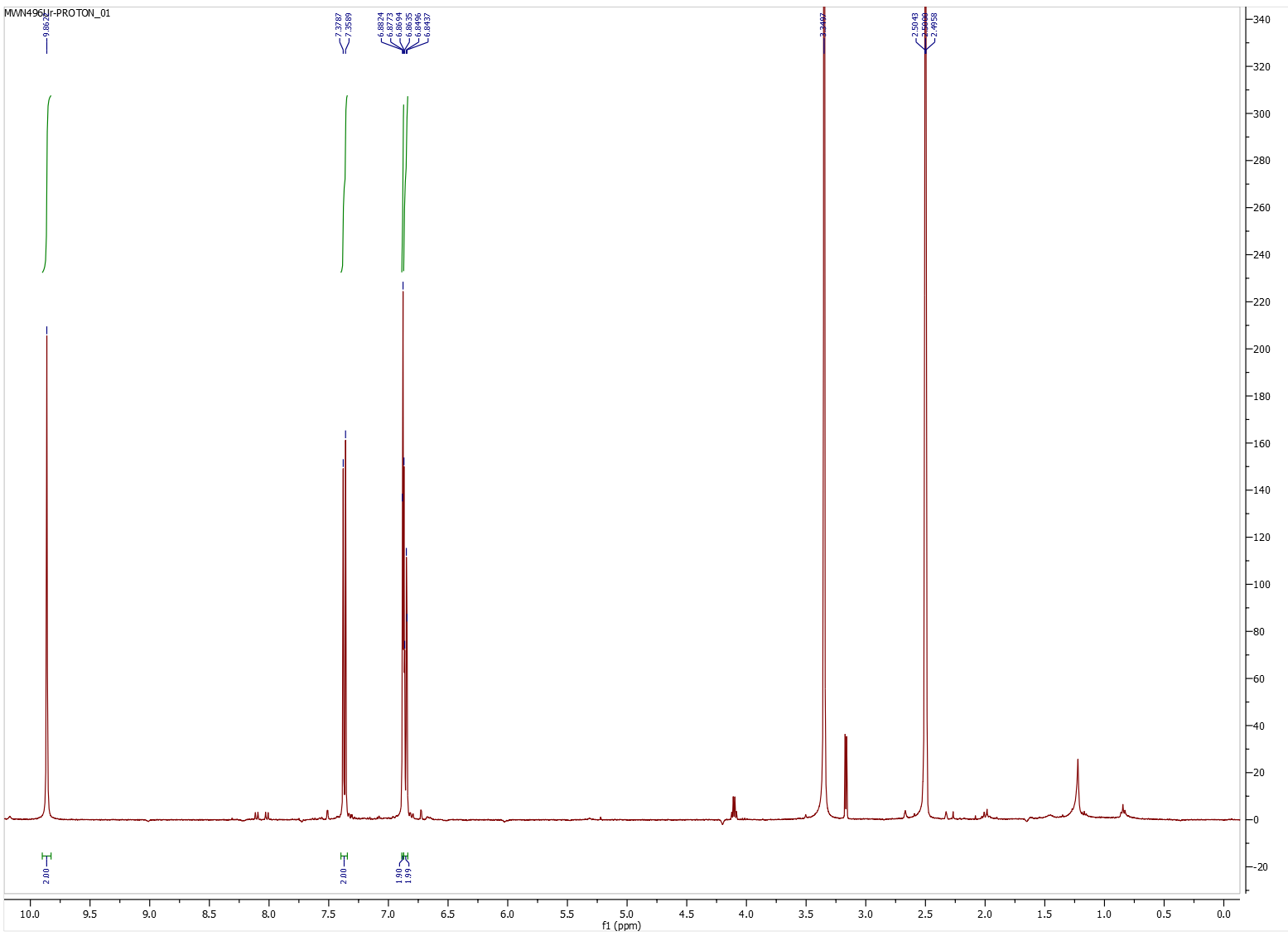

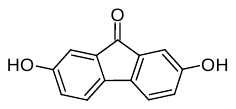

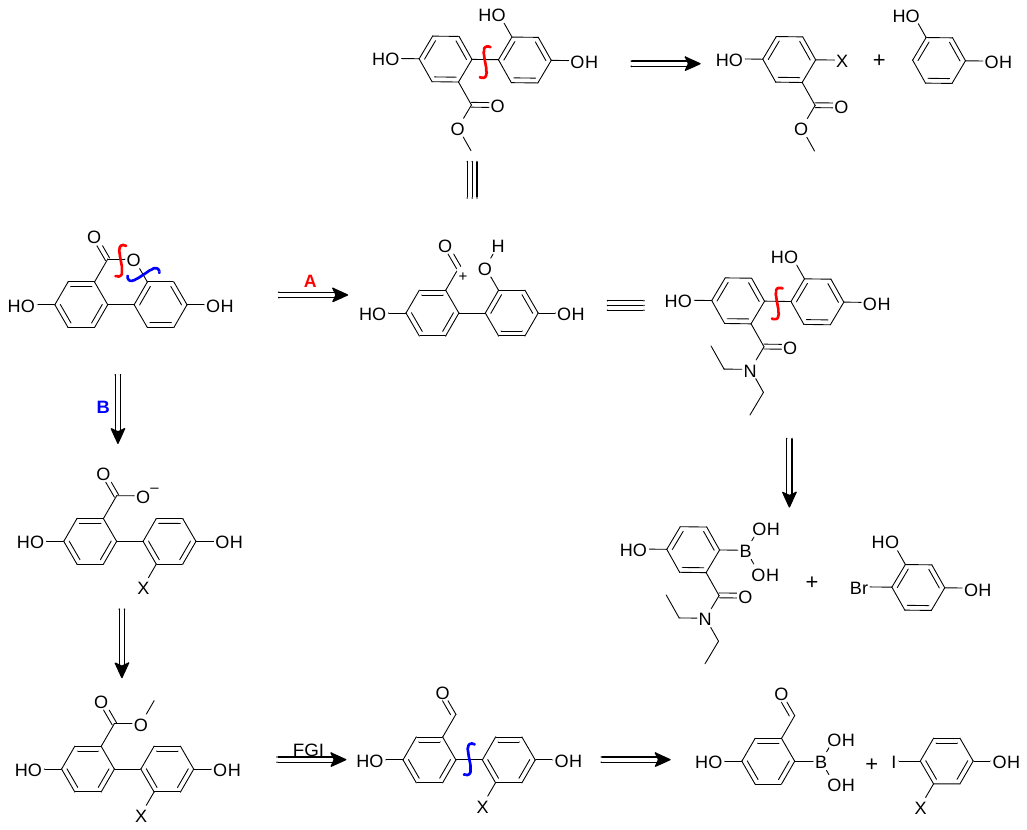

**Figure S1.** Common retrosynthetic strategies for UA.

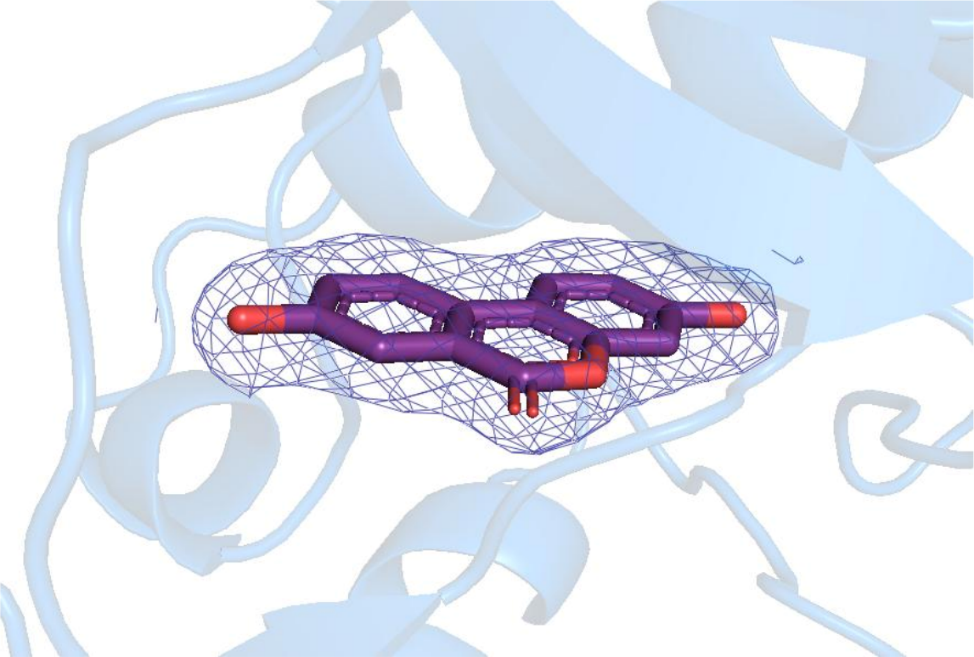

**Figure S2.** Electron density for UA in the DYRK1A/UA co-crystal structure. 2Fo-Fc electron density map contoured at +1.0σ (violet mesh) around UA (violet sticks; oxygen atoms in red) bound in the ATP-binding site of the DYRK1A kinase domain (blue cartoon), shown for chain A in the asymmetric unit (PDB ID: 35BW). The density covers the whole ligand and clearly defines its planar tricyclic scaffold and both hydroxyl groups. Figure was prepared with PyMOL 2.5.7.^14^

**Table S1.** Data collection and refinement statistics.

|  | **DYRK1A UROLITHIN A** |
| --- | --- |
| **PDB ID** | 35BW |
| **Wavelength** | 0.91840 |
| **Resolution range** | 48.68 - 2.13 (2.17 – 2.13) |
| **Space group** | P 21 21 21 |
| **Unit cell** | 88.76 89.85 229.29 (Å)  90 90 90 (°) |
| **Total reflections** | 601613 (30327) |
| **Unique reflections** | 103247 (5070) |
| **Multiplicity** | 5.8 (6.0) |
| **Completeness (%)** | 99.8 (91.7) |
| **Mean I/sigma(I)** | 9.1 (1.4) |
| **Wilson B-factor** | 33.0 |
| **R-merge** | 0.114 (1.158) |
| **R-meas** | 0.139 (1.407) |
| **CC1/2** | 0.996 (0.428) |
| **Reflections used in refinement** | 103354 |
| **Reflections used for R-free** | 5115 |
| **R-work** | 0.196 |
| **R-free** | 0.238 |
| **Overall number of atoms** | 11805 |
| **In macromolecules** | 10745 |
| **In ligands** | 72 |
| **In waters** | 988 |
| **Protein residues** | 345 (A), 342 (B), 339 (C), 336 (D) |
| **RMSD (bonds)** | 0.003 |
| **RMSD (angles)** | 0.639 |
| **Ramachandran favored (%)** | 96.92 |
| **Ramachandran allowed (%)** | 3.08 |
| **Ramachandran outliers (%)** | 0.00 |
| **Rotamer outliers (%)** | 0.00 |
| **Clashscore** | 1.75 |
| **Average B-factor** | 50.0 |

(13) *Grade2*; Global Phasing Ltd, Cambridge, United Kingdom: 2021. (accessed.

(14) *The PyMOL Molecular Graphics System*; Schrödinger, LLC.
